# Plant functional defects experienced upon growth under Per-/Poly-fluoroalkyl substances (PFAS) conditions

**DOI:** 10.64898/2026.08.14.743998

**Authors:** Jenson Lim, Nicola McKirdy

## Abstract

Per- and polyfluoroalkyl substances (PFAS) pose significant environmental risks, yet their impact on food crops like legumes remain insufficiently understood. This study investigated the developmental and physiological responses of hydroponically grown mung bean (*Vigna radiata*) to varying concentrations of perfluorooctanoic acid (PFOA) and perfluorooctanesulfonic acid (PFOS). High concentrations (1 mM) of PFOA severely impaired early plant development, significantly delaying seed germination, reducing leaf emergence, and suppressing root hair formation compared to PFOS and controls. Over a narrower concentration range (5–500 µM), both compounds caused transient growth stunting at early timepoints (48 h), though plants exhibited partial recovery over time. High-dose exposure (500 µM) significantly decreased seedling wet weights, leaf area, and leaf biomass without affecting dry weights, indicating disrupted water retention and homeostasis rather than reduced biomass accumulation. Spectrophotometric analysis revealed a dose- and compound-dependent effect on photosynthesis, with low-dose PFOA (5 µM) significantly increasing leaf chlorophyll absorbance. Furthermore, quantification of callose deposition revealed that high-dose PFOA (500 µM) and medium-dose PFOS (50 µM) raised baseline immune stress responses, which were not further elevated by subsequent flagellin-22 (flg22) challenge, suggesting a contaminant-induced immune priming mechanism. These findings highlight distinct, chemical-specific toxicological impact of PFAS on legume growth, water dynamics, and defence priming, underscoring critical implications for agricultural productivity and food safety.

## INTRODUCTION

Per- and polyfluoroalkyl substances (PFAS) are a diverse class of synthetic organofluorine compounds containing strong carbon-fluorine bonds, rendering them highly resistant to environmental degradation and thus known as “forever chemicals.” They exhibit thermal and chemical stability, oil-repellency, and surfactant properties which led to their widespread application in various industrial and consumer products (**Evich et al, 2025**). However, their presence in the environment, coming from manufacturing processes and effluent discharge, poses significant ecological and human health concerns due to their toxicity and ability to bioaccumulate in various matrices (including soil and water) and organisms (**Oviedo-Vargas et al, 2025; Caniglia et al, 2022; Dimitrakopoulou et al., 2024**).

Under the umbrella of PFAS, perfluorooctanoic acid (PFOA) and perfluorooctanesulfonic acid (PFOS) well-characterised PFAS due to their presence in waterbodies, largely attributable to industrial emission, dispersion, spills, and disposal of waste and wastewater. The use of firefighting foams has also been implicated in this too (**Wee & Aris, 2023**). PFAS uptake, accumulation and effects on terrestrial flora, particularly edible crops, is understudied and a critical knowledge gap, as they serve as pathways for human exposure (**Li et al, 2022; Ecke et al., 2025**). For example, PFAS is present in biosolids, which are often applied as fertilizers, thereby introducing PFAS into agricultural soils (**Biswas et al., 2025**). Therefore, this necessitates a comprehensive understanding of how PFAS interacts with, and affects edible crops, as this represents a critical pathway for human exposure. Due to the nature of plant field studies with PFAS introduced to growing seedlings, potential effects on germination and early growth are missed (**Costello and Lee, 2024**). Furthermore, the field studies involve soil which affect PFAS availability due to the nature of soil sorption.

To study the impact of PFAS on seed germination and its subsequent growth, mung bean (*Vigna radiata*) was adopted, and grown hydroponically. As a crop, mung beans are relatively fast growing, reaching maturity in 75 to 90 days. They are also tolerant to environmental stresses, such as high temperatures, drought, and salinity. Legumes, which includes mung beans, are cultivated worldwide for its nutritional and dietary value. However, its use in Western society has reduced over the past centuries. Yet, there is renewed interest in legumes due to its sustainability and nutritional value as a food source as well as fodder (**Ferreira et al, 2021; Li et al., 2017**). Legumes provide a cheap plant-based protein source, is more environmentally friendly and protects natures’ biodiversity (**Everwand et al, 2017**). All these properties make mung beans a desirable target for cultivation. The increasing concern over PFAS contamination underscores the need to investigate their impact in agricultural crops, which is vital for global food security.

Currently, there is limited literature that examines the effects of PFAS on mung beans, a plant species that is staple in Asian cuisine and with increasing consumption worldwide as they are a good and inexpensive source of antioxidants and nutrients (**Peñas et al, 2010; Han et al., 2022; Kapravelou et al., 2020**). Understanding the interaction between PFAS and mung bean plants is crucial for assessing potential risks to agricultural productivity and human health through dietary exposure (**Khalil et al., 2020**). Therefore, this research aims to elucidate the effects of varying PFOA and PFOS concentrations on mung bean growth and development.

## MATERIALS AND METHODS

### Reagents

Mung beans (*Vigna radiata,* Lotus Foods Trading Company Ltd., UK) was purchased from Amazon.co.uk. All key reagents, such as aniline blue (Fisher 10385450), ethanol, glycine, HCl and NaOH, perfluorooctanoic acid (PFOA, Acros 173960050), perfluorooctanesulfonic acid (PFOS, Aldrich 77283) were purchased from either Fisher Scientific or Sigma-Aldrich, UK in their purest form.

### *Vigna radiata* germination and growth

To understand the broad effects of PFAS on plant development, *Vigna radiata* seeds were exposed to a wide range of PFOA- and PFOS-containing solutions, two of the most commonly used, and widely studied PFAS. Standard 1 pound jam jars 11 cm (height) x 7 cm (diameter) containing 1 folded piece of blue roll (30 cm x 20 cm, Amazon Commercial) each were sterilised by autoclaving. 3-5 seeds were place into each jar and between 12-30 seeds per sample/control were used per experiment. Each experiment was performed on 3 separate occasions. As the main impact of PFAS lies between 1μM and 1mM, focus was initially placed within that range before narrowing to 5 – 500 μM. *Vigna radiata* seeds were exposed to 5 – 500 μM PFOA- and PFOS-containing solutions and tracked over time for hypocotyl and leaf growth. PFOA or PFOS was prepared in 30 ml of distilled water and added to the jars. 6 – 10 seeds were added per jar, lidded with aluminium foil and incubated at 25 °C in complete darkness.

After a growing for up to 29 days, some seedlings were taken out, and the hypocotyls were measured with a ruler from the base of the first leaves to the top of the root system. Leaves were counted, scanned and the area of each leaf was measured using the ImageJ software (National Institutes of Health). At least 6 and up to 9 leaves from each condition were measured across three independent experiments. Number of root hairs were also counted visually. All were blind measured and counted.

### Quantitative determination of extracted chlorophyll and callose

To ascertain further effects that PFAS might have on leaf growth and development, we explored the biochemical changes to mung bean plants grown in the presence of PFAS. We explored if chlorophyll levels were affected by PFAS as chlorophyll is critical for photosynthesis, the process of substance and energy circulation in plants (**Pavlovi**ć **et al 2014**). These levels of chlorophyll within plants decrease when facing environmental stressors such as heat and drought (**Hannahchi et al 2022**). Prior to chlorophyll extraction, plants were grown to a fixed timepoint and their leaves weighed. Chlorophyll and callose extraction method was adapted from **Schenk and Schikora (2015)**. Seedlings were transferred to individual jars containing clay pebbles (02-050-105, VitaLink, UK) and tap water and allowed to grow for 29 days. Two to three true leaves were removed, weighed and where appropriate, suspended in 1 μM flagellin peptide 22 (flg22; GenScript RP19986) in hydroponic water (VitaLink HydroMax Grow) for 24h. Leaves were later homogenised in a fixed volume of 1:3 glacial acetic acid:ethanol (GAA:EtOH) using a 1.5 ml homogeniser. Destained leaf fragments were centrifuged for 20 min at 12000 g in a microcentrifuge and GAA:EtOH was removed from destained leaves by careful pipetting and measured in a plate reader (Molecular Devices VersaMax plate reader) at 410 nm and 660 nm, for chlorophyll.

Next, pellets of leaf fragments were dissolved in 300–350 μl DMSO, sealed with parafilm tape and autoclaved at 121°C for 20 min, 20 psi. Once tubes were cooled to room temperature, they were centrifuged at 12000 g for 5 min before transferring supernatants into fresh tubes. 50 μl of supernatant was supplemented with 100 μl 1 M NaOH and 0.6 ml loading mixture (containing 200 μl 0.1% (w/v) aniline blue in water, 295 μl 1 M glycine/NaOH (pH 9.5) and 105 μl 1 M HCl), mixed vigorously, then incubate in a water bath at 50°C for 20 min. Once cooled to room temperature (approximately 30 min), 100 μl was aliquoted to 3 wells in a 96-well plate black opaque plate and read in a microplate reader with fluorescence capabilities at 365 nm/500-550 nm excitation/emission wavelengths. These are the readings for callose, and they were corrected with leaf weights (**Kohler et al, 2000**).

### Statistical analyses

ANOVA or non-linear least square fit regression were used, along with Tukey’s and sum-of-square F-test multiple comparisons test, respectively, to assess the effects of PFAS for all experimental endpoints. EC_50_ was determined using non-linear fit (variable slope). All analyses were performed in GraphPad Prism 9.4.0., San Diego, California USA, www.graphpad.com. Samples sizes can be found within the respective methods sections.

## RESULTS

### PFOA, not PFOS has toxic and developmental effects on *Vigna radiata* at high doses

Seed germination (**Figure 1**) and its subsequent root & leaf formation (**Figure 2A – C**) were tracked over time and it was found that PFOA was more toxic to seeds compared to PFOS, at high concentrations (1 mM), as evidenced by lower and slower seed germination (**Figure 1**). Furthermore, from the seeds that germinated, while there was a decrease in the number of root hairs (**Figure 2A**) and hypocotyl length (**Figure 2B**) at the highest dose (1 mM) of both treatments, PFOA-treated seed exhibited more significant effects, compared to PFOS (**Figure 2A**; 6.00 ± 4.19 c.f. 9.57 ± 4.98 [control]; *p* = 0.0235) (**Figure 2B**, 29.40 ± 16.18 c.f. 67.33 ± 49.38 [control]; *p* = 0.0198). Interestingly, seeds treated with PFOA at 1nM, had a small, though insignificant decrease in the number of root hairs and length of roots. Unsurprisingly, leaf formation was affected too, with seedlings treated with the highest dose of PFOA having fewer leaves (**Figure 2C**, 1 mM; 4.00 ± 5.29 c.f. 17.00 ± 2.65 [control]; *p* = 0.0082). At 1nM concentration, PFOS reduced the number of leaves too, albeit not statistically significant. This suggests PFOA can cause major effects on mung bean seedlings occur early in the plant’s life.

**Figure 1:**
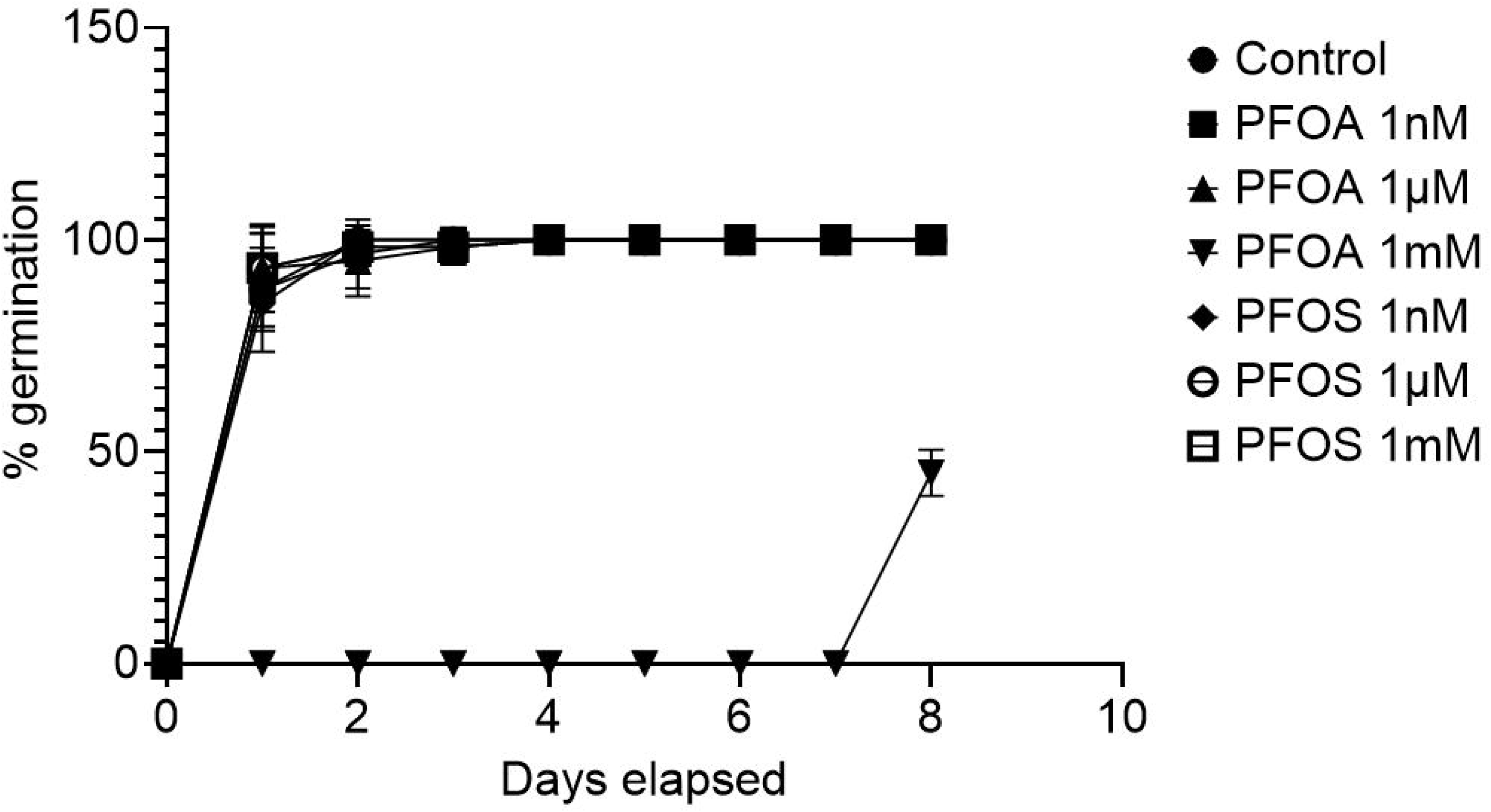
High dose PFOA affects *Vigna radiata* seed germination. Mung bean seeds were allowed to germinate on paper towels soaked with the appropriate PFAS and tracked for 8 days as described in the Materials and Methods. Results were based on the total of two independent experiments (n = 20).

**Figure 2:**
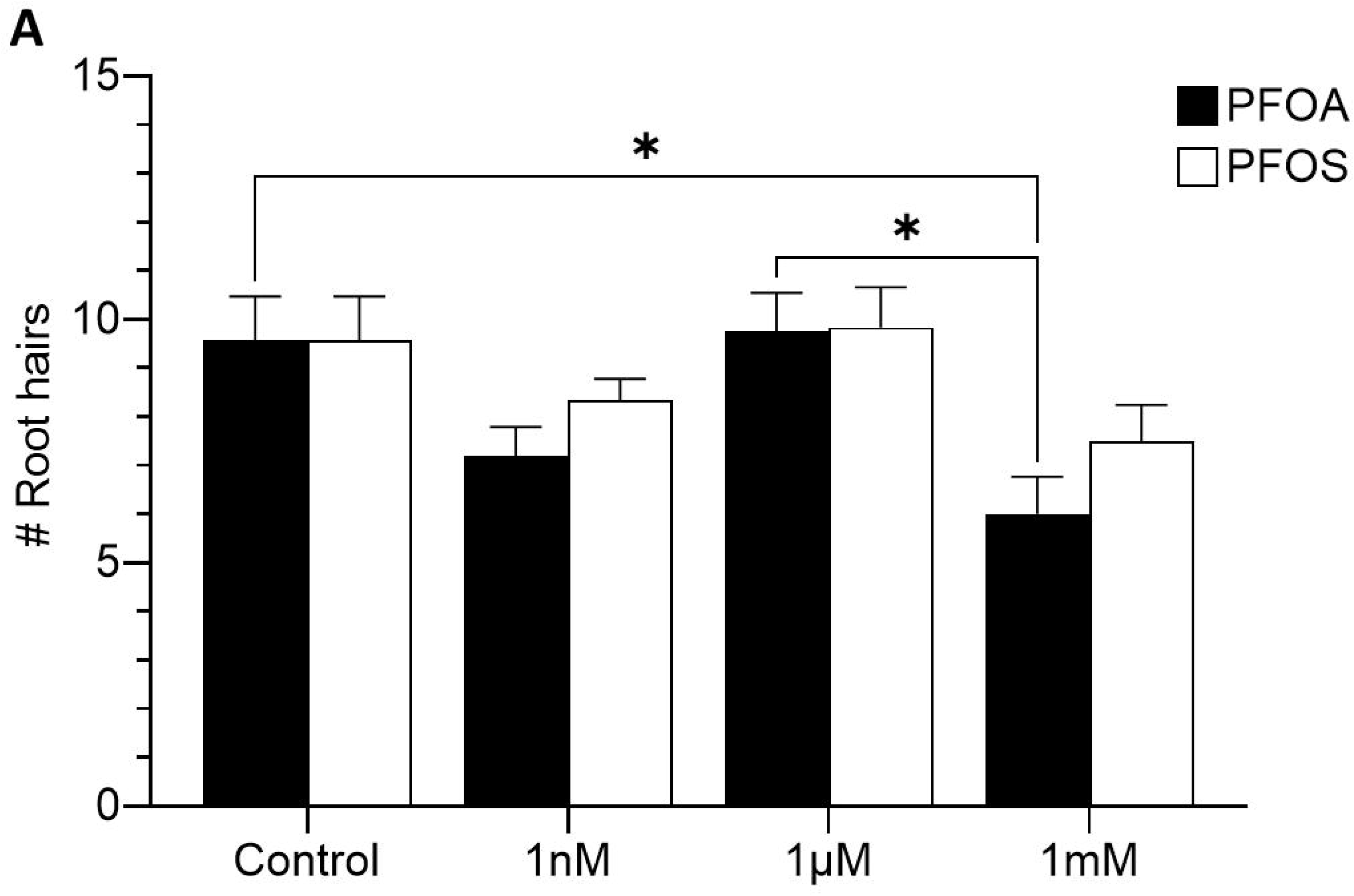
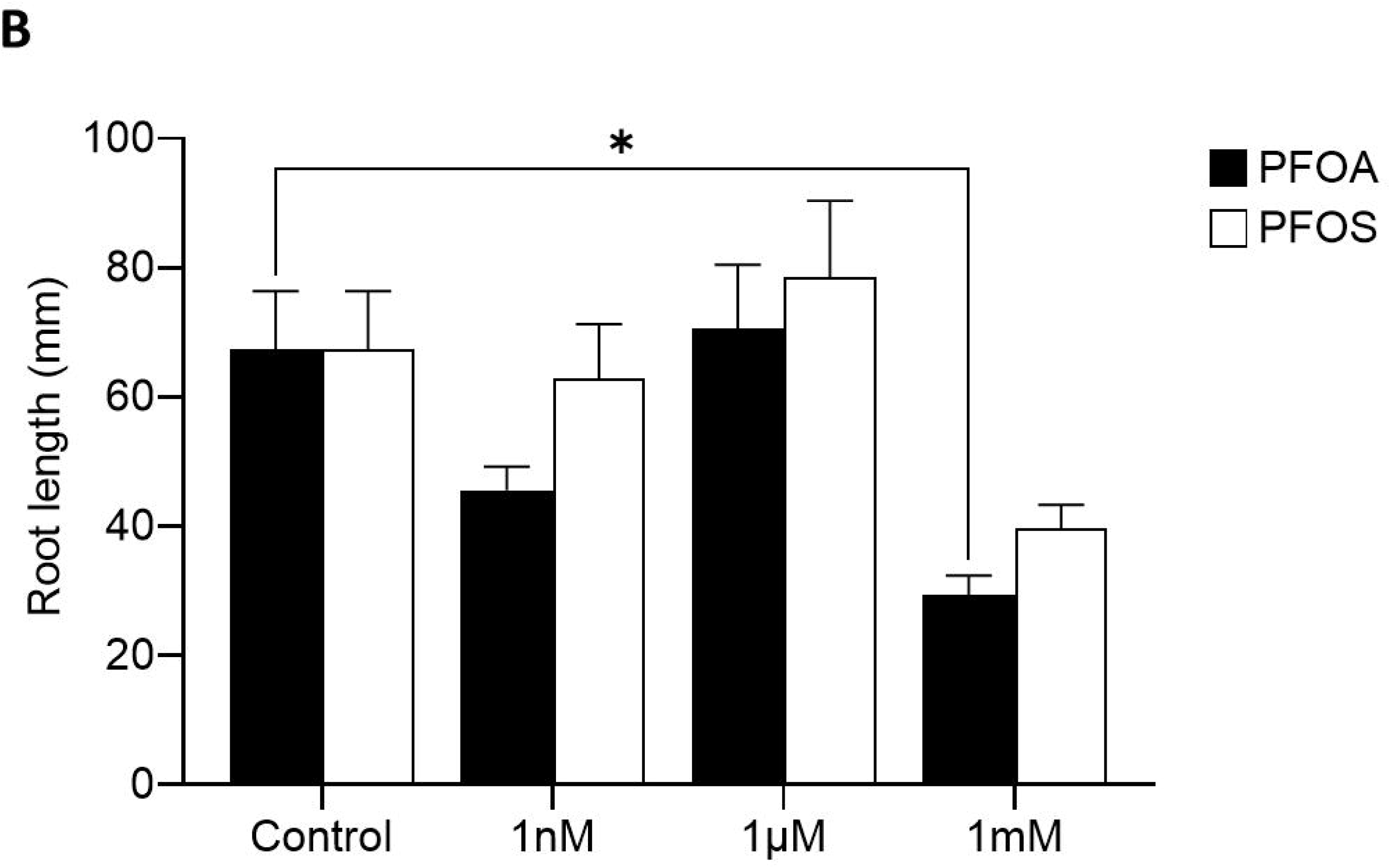
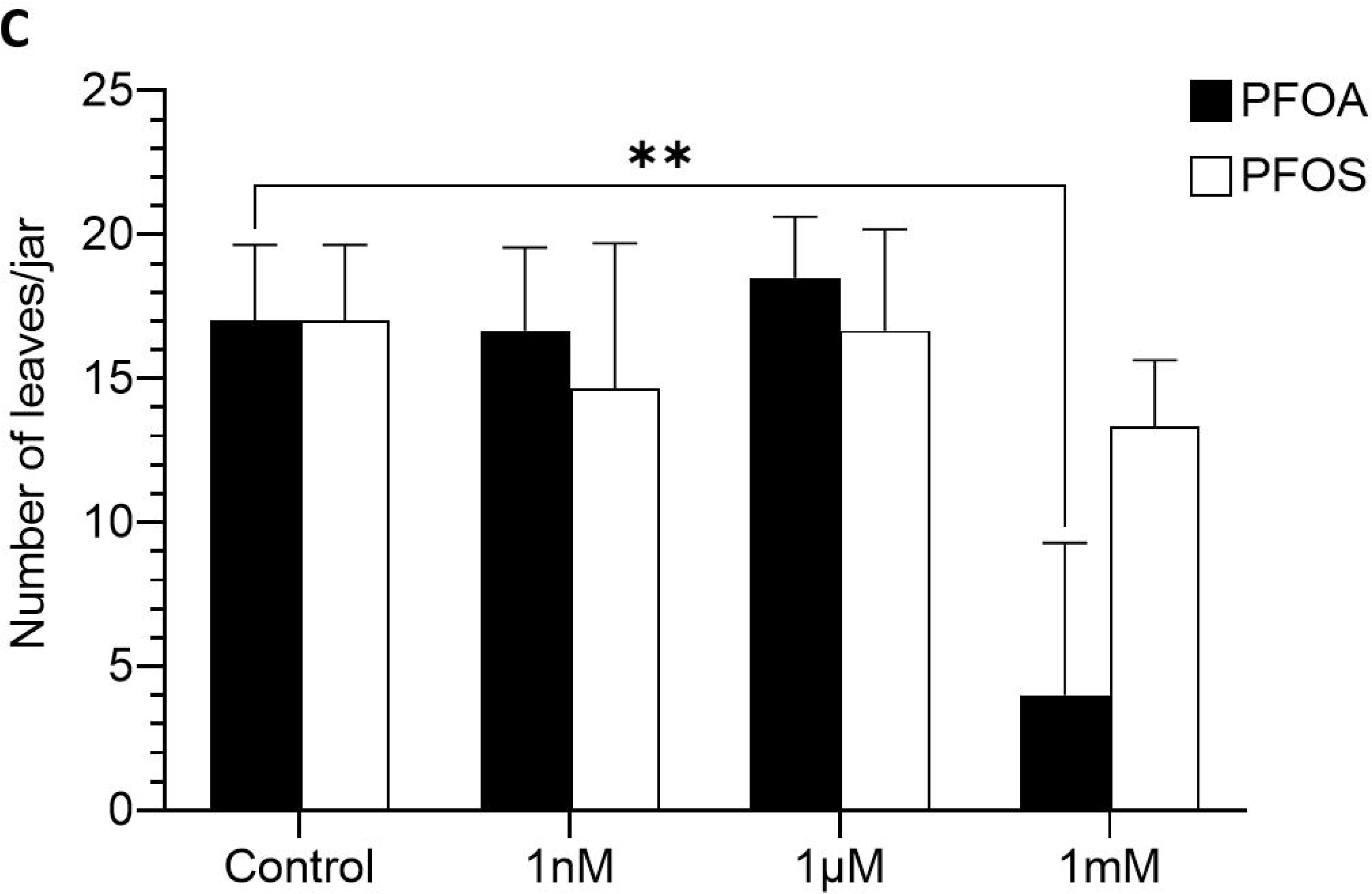
High dose PFOA affects *Vigna radiata* seedling development. Mung bean seeds were germinated at 25°C and grown with the appropriate PFAS for 21 days, as described in the Materials and Methods. Individual seedlings were counted for root hairs (**A**), root lengths (**B**) and number of leaves per jar (**C**). Statistical significance compared to untreated plants was determined by two-way ANOVA, and a Tukey’s multiple comparisons test. (**) p ≤ 0.01 (*), p ≤ 0.05, otherwise, not significant. Results are expressed as the mean ± SEM of at least three independent experiments (n = 30)

### PFAS affects overall seedling growth

As the main impact of PFAS lies between 1μM and 1mM, focus was placed within that range. *Vigna radiata* seeds were exposed to 5 – 500μM PFOA- and PFOS-containing solutions and tracked over time for hypocotyl and leaf growth. It was found that, in general. Overall, both PFOA and PFOS led to slightly stunted seedling growth, though it was more significant for PFOS compared to PFOA, at low (5 μM; 35.92 ± 11.55 mm c.f. 61.92 ± 14.32 mm [control]) and high doses (500 μM; 27.92 ± 16.14 mm c.f. 61.92 ± 14.32 mm [control]; **Figure 3A**). Interestingly, this phenomenon was observed at 48 h, not 60 h, when both PFOA and PFOS stunts growth at all concentrations, compared to its untreated control (**Figure 3B**). This suggests that while the plants may be stunted by PFOS initially, this is rectified over time, likely through a compensation mechanism.

**Figure 3:**
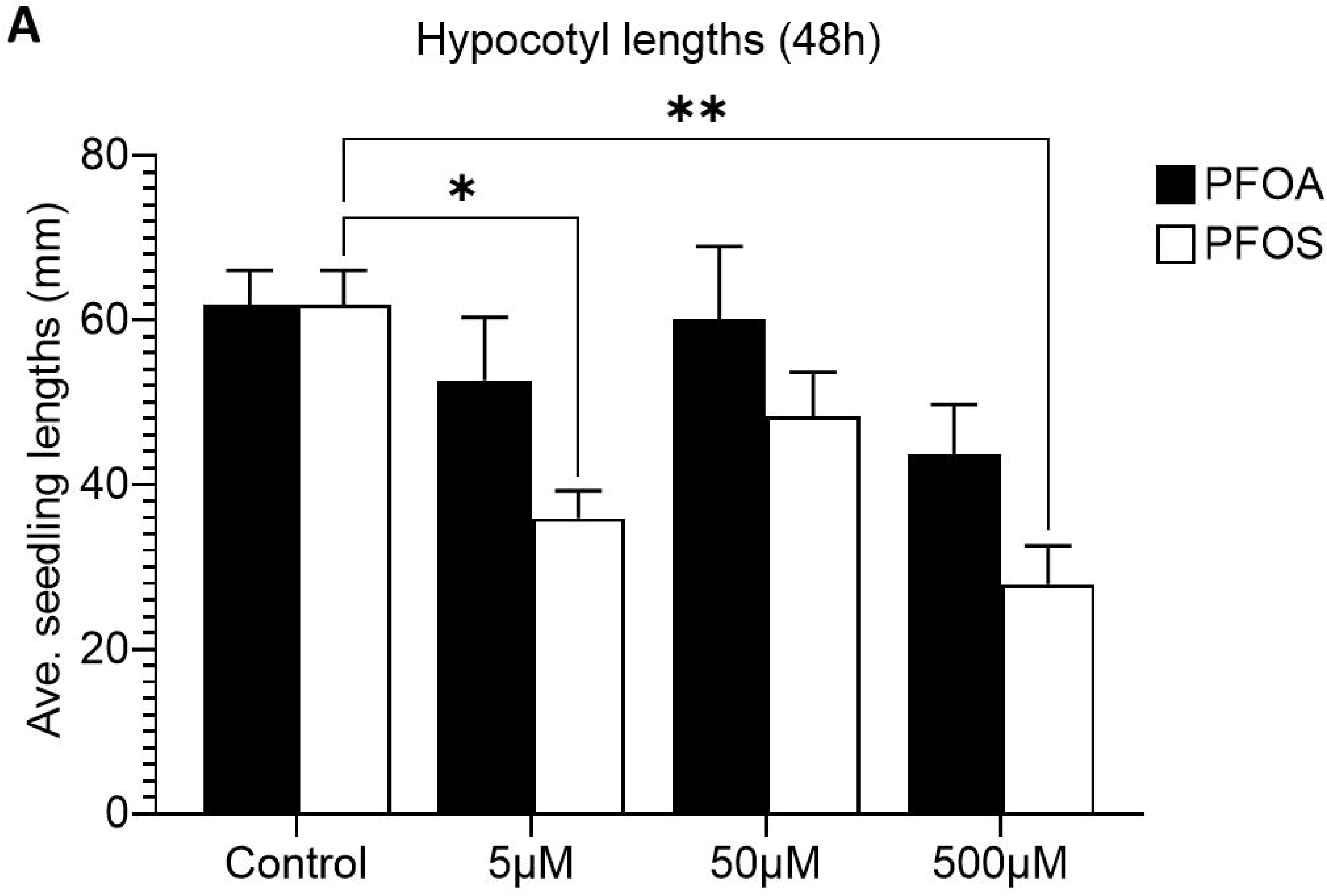
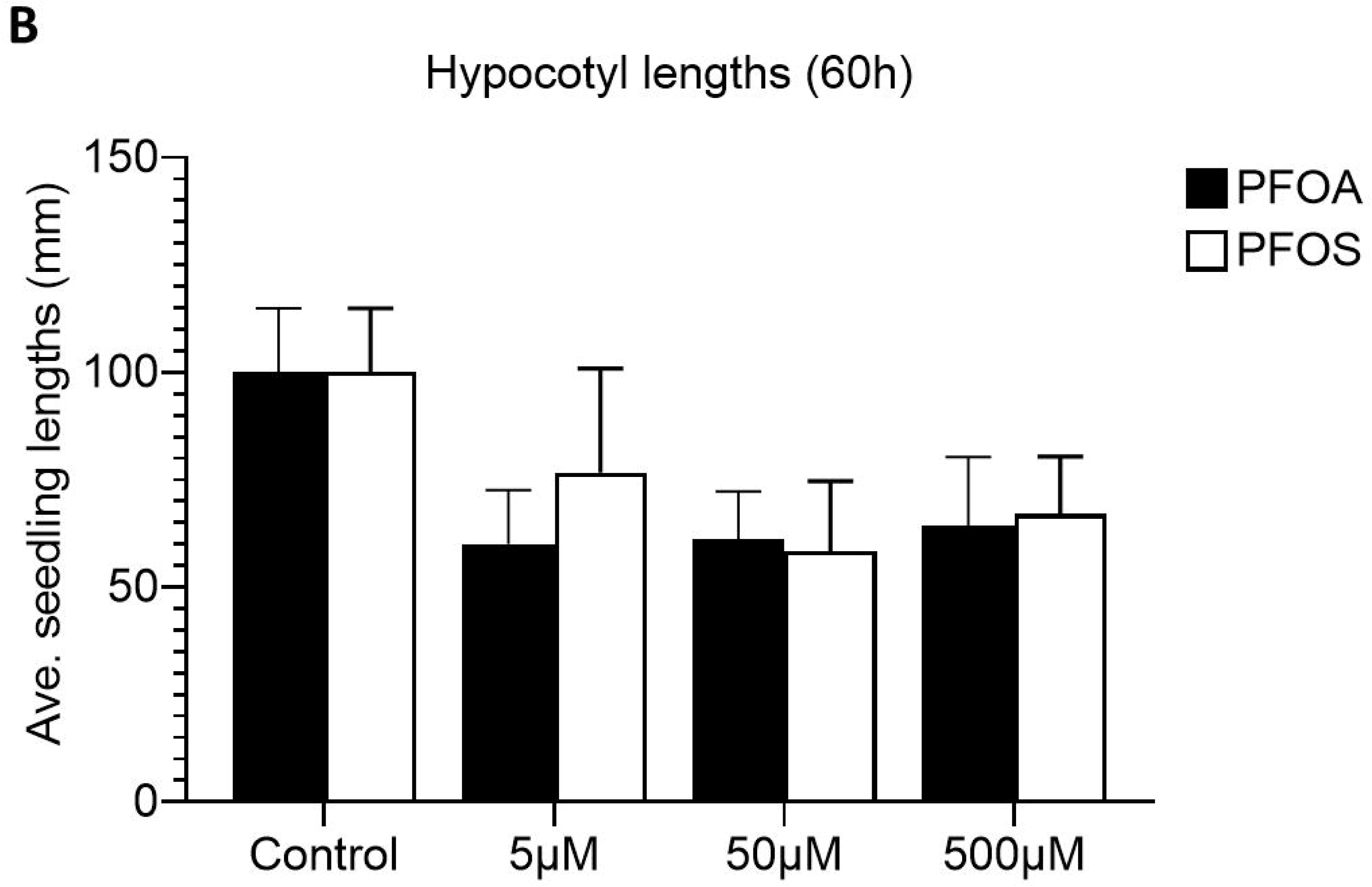
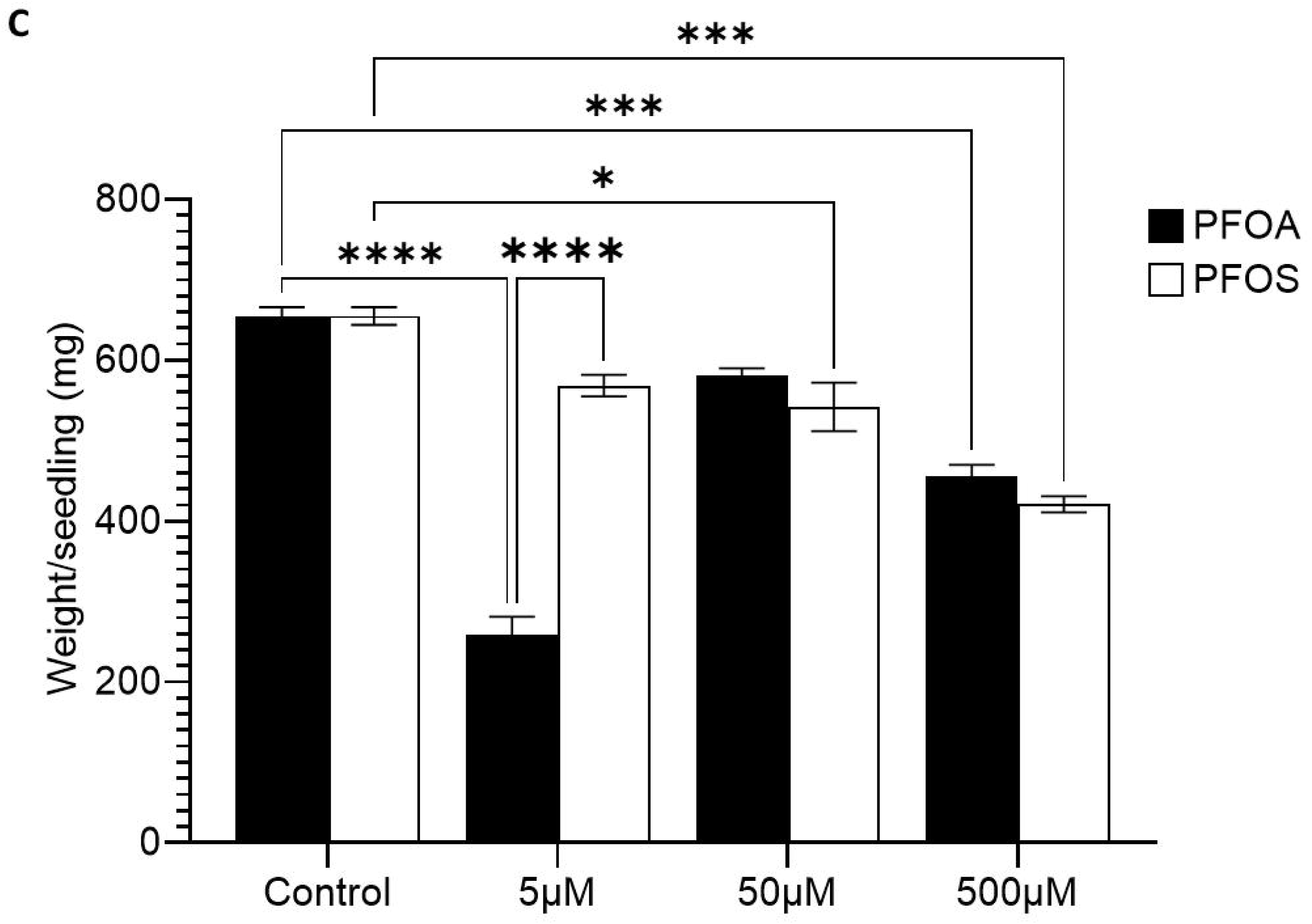
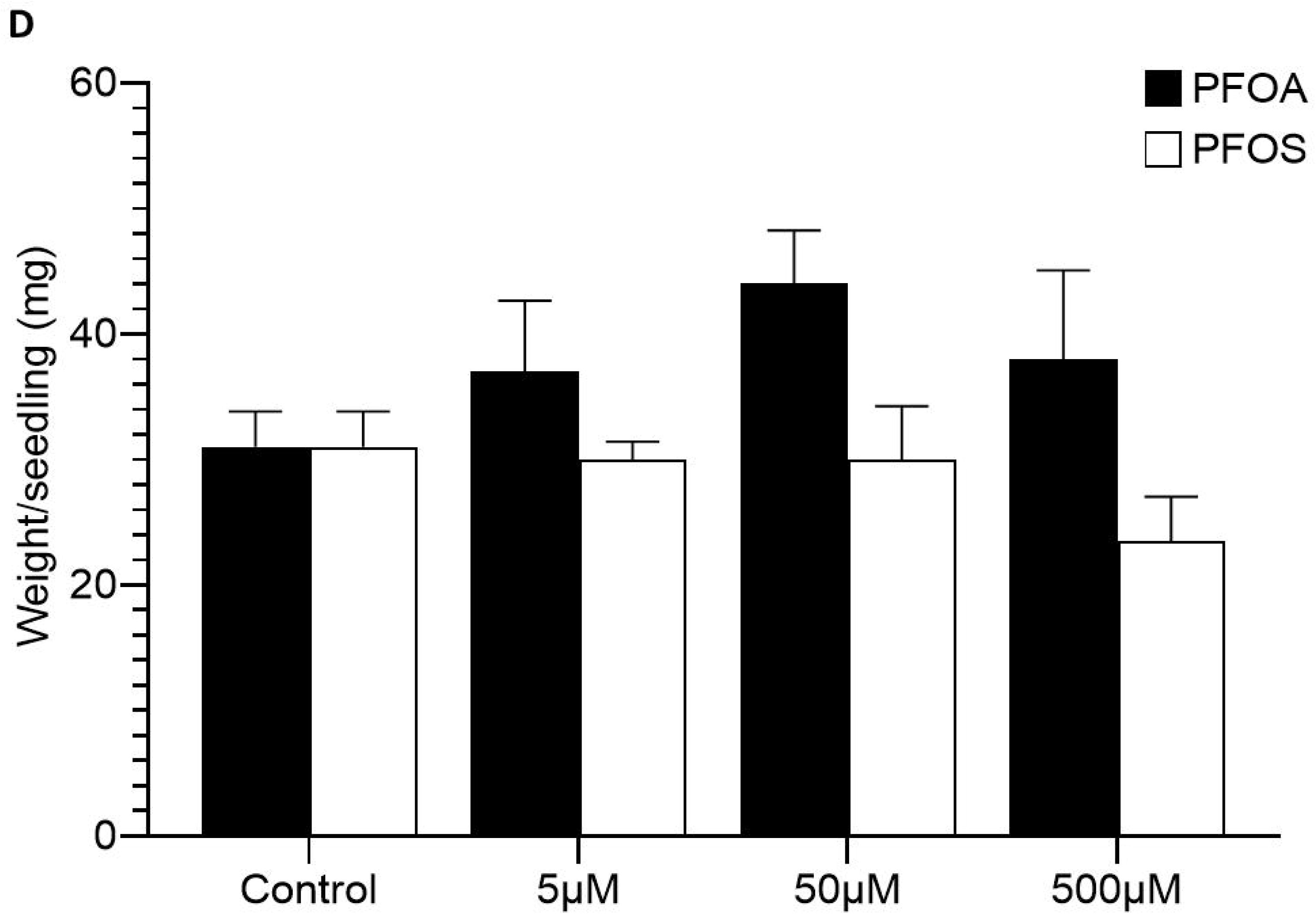
PFAS affects hypocotyl and leaf growth. Mung bean *(Vigna radiata)* seeds were germinated and allowed to grow for 48 h (**A**), 60 h (**B**) or 28 days (**C, D**) before hypocotyls (**A, B**) or entire seedlings were weighed (**C**), dried and weighed again (**D**), as described in the Materials and Methods. Statistical significance compared to untreated plants was determined by two-way ANOVA, and a Tukey’s multiple comparisons test. (****) p ≤ 0.0001, (***) p ≤ 0.001, (**) p ≤ 0.01 (*), p ≤ 0.05, otherwise, not significant. Results are expressed as the mean ± SEM of at least two independent experiments (n = 9-12 leaves from different plants/condition)

However, after 28 days, when the weights of seedlings were determined, it was revealed that while low doses (5 μM) of PFOA had a negative impact on wet weights, at higher doses (500 μM) of both PFOA and PFOS led to decreased wet weights compared to the untreated controls (**Figures 3C**). This suggests a dose- and chemical-dependent response on the wet weight of mung beans. On the other hand, dry weights from seedlings grown in both PFOA and PFOS has slight decrease in weight compared to the controls, albeit not at significant levels (**Figure 3D**). At low and medium doses, results were less clear with seedlings having lower wet weight when grown in 5 μM PFOA (258.00 ± 32.53 mg c.f. 655.00 ± 15.56 mg [control]) or 50 μM PFOS (542.00 ± 42.43 mg c.f. 655.00 ± 15.56 mg [control]).

### PFAS affects leaf development and chlorophyll levels

To ascertain further effects that PFAS might have on mung bean leaf growth and development, seeds were germinated and allowed to grow for 29 days in PFOA or PFOS to display leaves of different sizes (**Figure 4A**). In general, true leaves from seeds grown in the presence of PFOA and PFOS were smaller than the controls. That is certainly true at a higher dose of 500 μM of chemical. (PFOA, 43854.83 ± 5706.99 c.f. 72531.78 ± 4224.91 [control]; *p* = 0.004; PFOS, 41310.5 ± 3695.04 c.f. 72531.78 ± 4224.91 [control]; p = 0.001). Interestingly, low (5μM) dose of PFOA drastically decreased leaf sizes (31239.5 ± 3360.53 c.f. 72531.78 ± 4224.91 [control]; *p* ≤ 0.0001). Furthermore, this result is also complemented by the corresponding leaf weights (**Figure 4B**) highlighting that the development of seedlings are negatively affected by a narrow range of PFAS.

**Figure 4:**
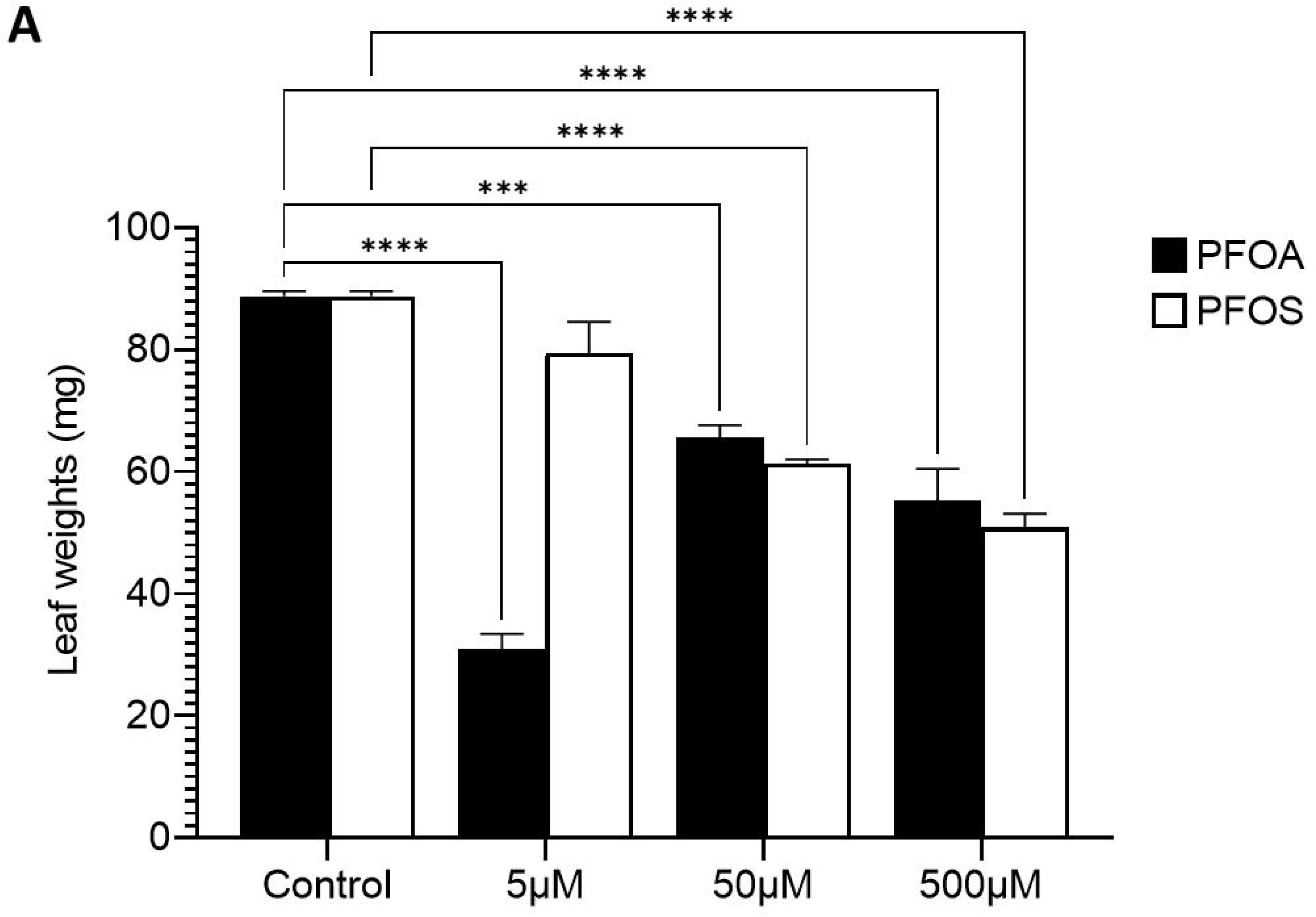
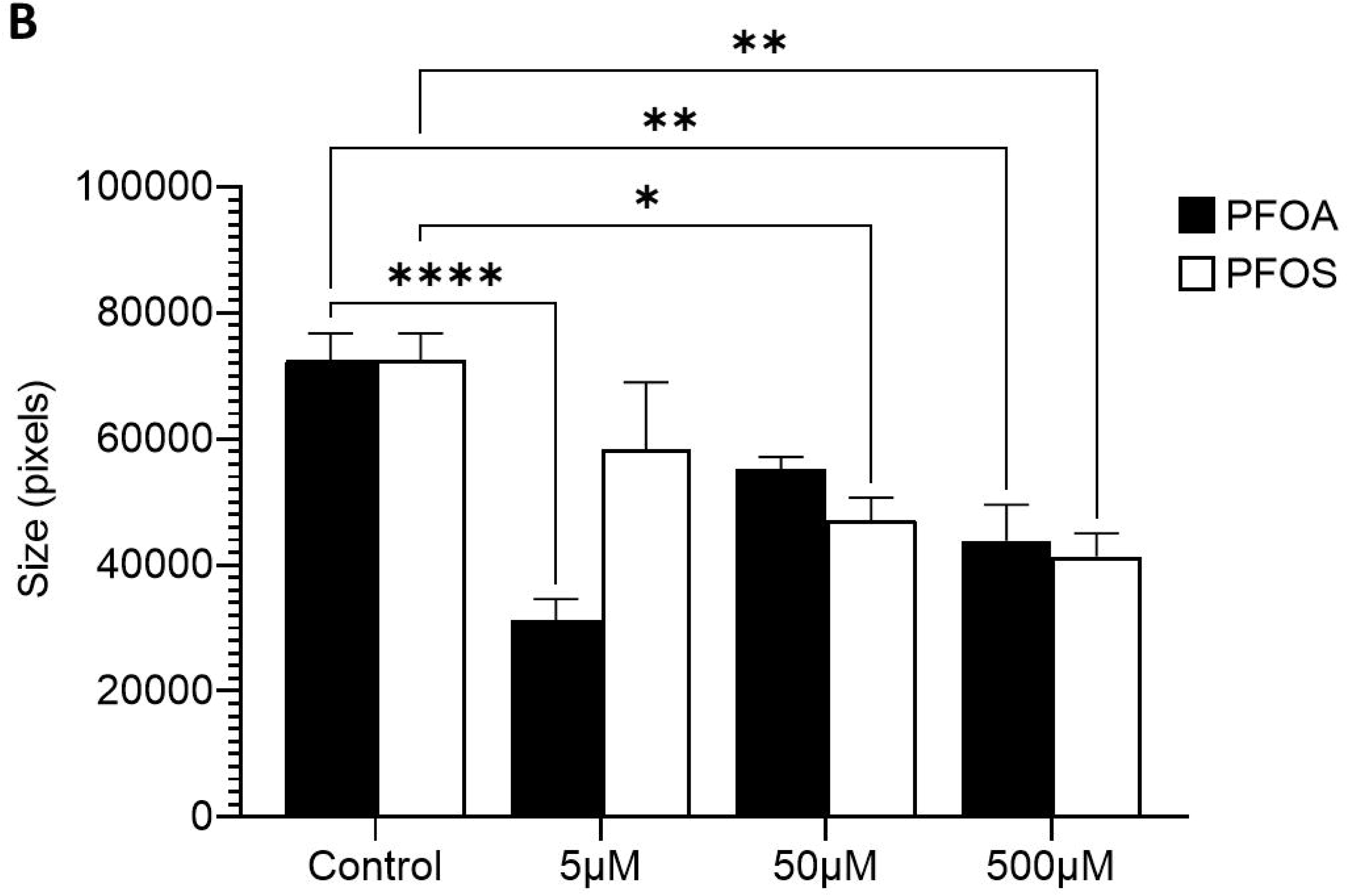
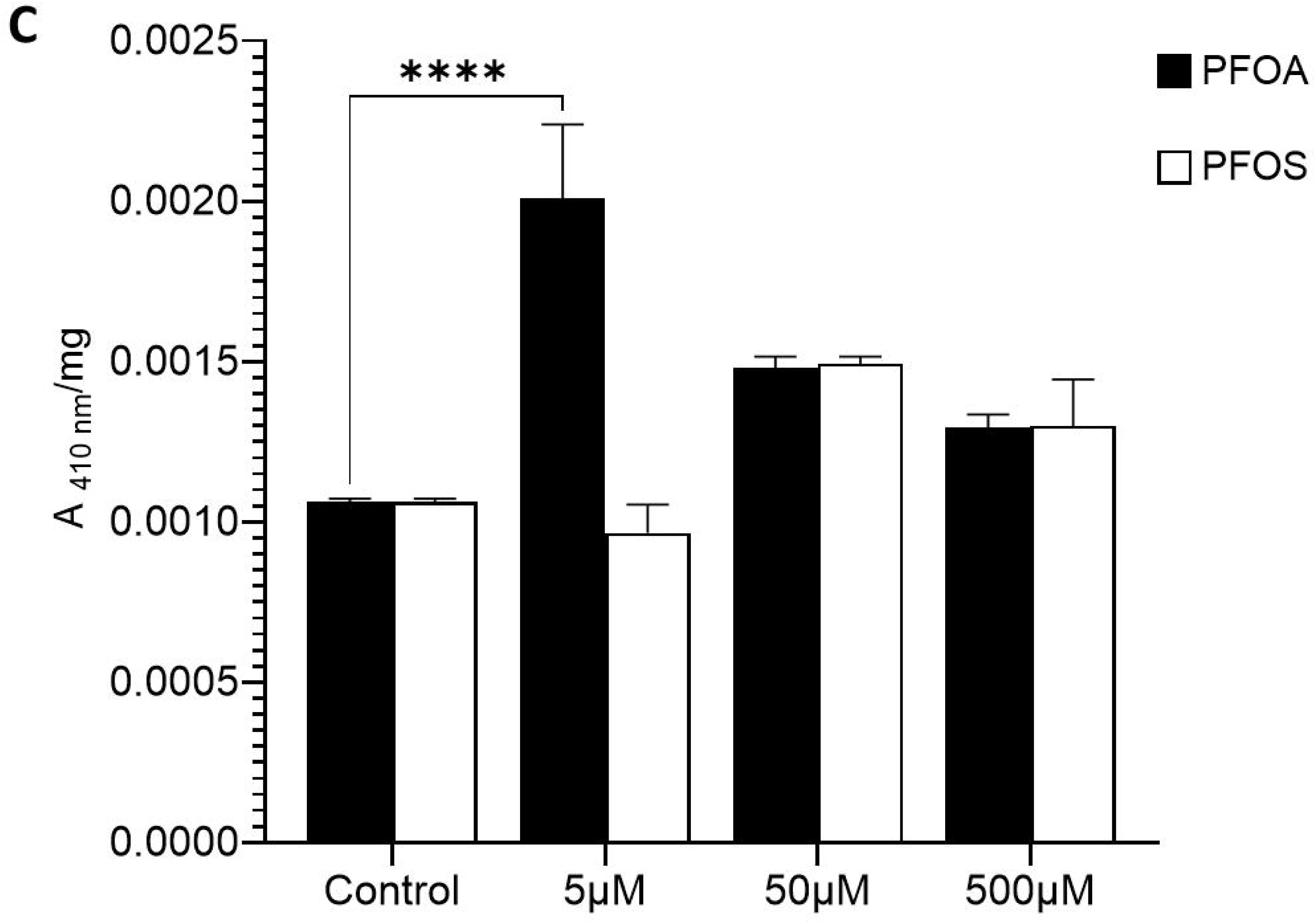
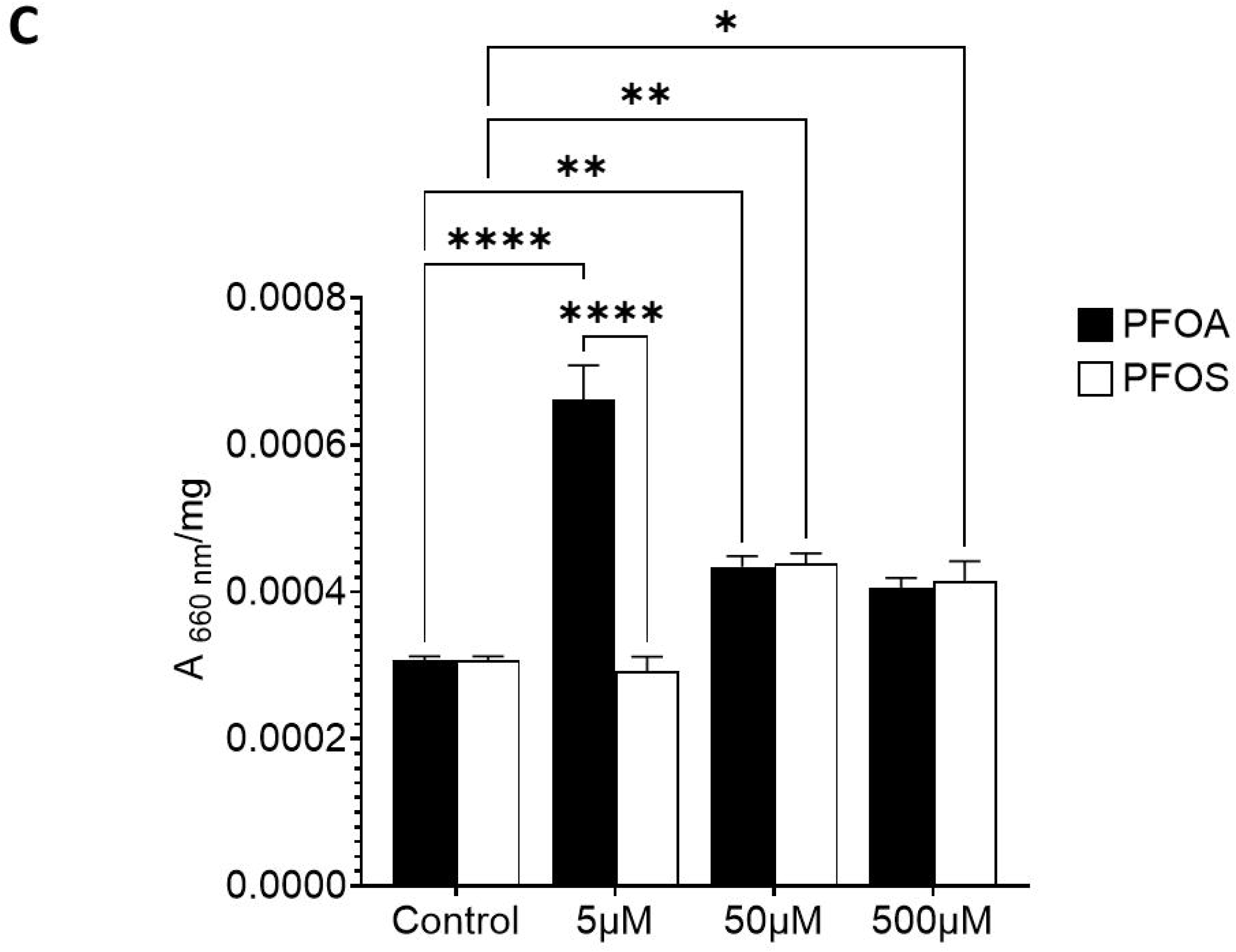
Chlorophyll and weight levels are altered by PFAS in *Vigna radiata* plants. Mung bean seeds were germinated and allowed to grow for **29** *(15/10/2021 – 12/11/2021)* days. Leaves were removed and either weighed **(A)** or scanned and measured for pixel size (**B**). Chlorophyll was extracted from leaves were and measured spectrophotometrically and normalised against the weight of leaves (**C**), as described in the Materials and Methods section. Significance was determined by two-way ANOVA, and a Tukey’s multiple comparisons test. (****) p ≤ 0.0001, (***) p ≤ 0.001, (**) p ≤ 0.01 (*), p ≤ 0.05. Results are expressed as the mean ± SD of at least two independent experiments (n = 8 leaves/condition).

At low dose, PFOA but not PFOS led to significantly lighter leaves (5 μM; 30.83 ± 5.19 c.f. 88.75 ± 1.71 [control]; p < 0.0001) in line with the previous result of lighter seedlings (wet weight; **Figure 3C**) and smaller leaves (**Figure 4A**). At medium doses (50 μM; PFOA, 65.63 ± 3.99; PFOS, 61.28 ± 1.48) and high (500 μM; PFOA, 55.20 ± 10.52; PFOS, 50.90 ± 4.38), there was a significant decrease in leaf weights, compared to the control (88.75 ± 1.71), in a dose dependent manner. As PFAS affect leaf weights and sizes, we wanted to determine if chlorophyll levels were also affected. First, chlorophyll was extracted from the true leaves of control seedlings and the absorption spectrum determined. While chlorophyll a and b could not be discriminated spectrophotometrically, possibly due to the sensitivity of the plate reader, we recorded the maxima of chlorophyll and determined them to be 410 nm and 660 nm which is expected of most green plants (**Supplementary Figure**). True leaves from seedlings grown in PFOA and PFOS were removed, chlorophyll extracted and measured spectrophotometrically, at the two maxima (**Figure 4C [410 nm], 4D [660 nm]**). Interestingly, while there was a modest increase in absorbance from seedlings grown in 50 and 500μM PFOA and PFOS, this highest increase in absorbance (410 nm and 660 nm) was found in extracts from seedlings grown in 5μM PFOA (0.0007 ± 0.00012 c.f. 0.0003 ± 0.00001 [control], p < 0.0001) but not PFOS (**Figure 4C, 4D**). This suggests that mung bean seeds grown in low dose PFOA, but not PFOS, could promote an increase in chlorophyll production in plants.

### PFOA, not PFOS induces a stress response in plants

One of the plant’s responses to stress is the accumulation of callose, which is detectable by aniline blue (**Chen and Kim, 2009; Schenk and Schikora, 2015**). Leaves from control plants were challenged with flagellin (flg22) to activate their stress responses. This produced callose which was detected by aniline blue (72.33 ± 2.69), when compared to untreated (60.66 ± 0.487; p < 0.0001) or aniline blue-free leaves (29.75 ± 0.21; p < 0.0001) (**Figure 5A**). Next, leaves from plants grown for 29 days in either PFOA or PFOS were challenged with Flg22 before tested for the presence of callose using aniline blue. It was found that there was only a modest increase in callose production in leaves from untreated (no PFAS) plants challenged with Flg22 (5.50 ± 0.21 [-Flg22] c.f. 7.27 ± 0.95 [+ Flg22]; p < 0.0001) (**Figure 5B, 5C**). However, there were significant increases of callose production for PFOA at high dose (500 μM; 10.19 ± 0.83; p < 0.0001) or PFOS at medium dose (50 μM; 7.93 ± 1.53; p = 0.0001) when compared to the untreated control (5.50 ± 0.21). This suggests that the stress response in plants is primed mainly by PFOA, and less so by PFOS, possibly to confer an elevated baseline level of protection from further antagonism.

**Figure 5:**
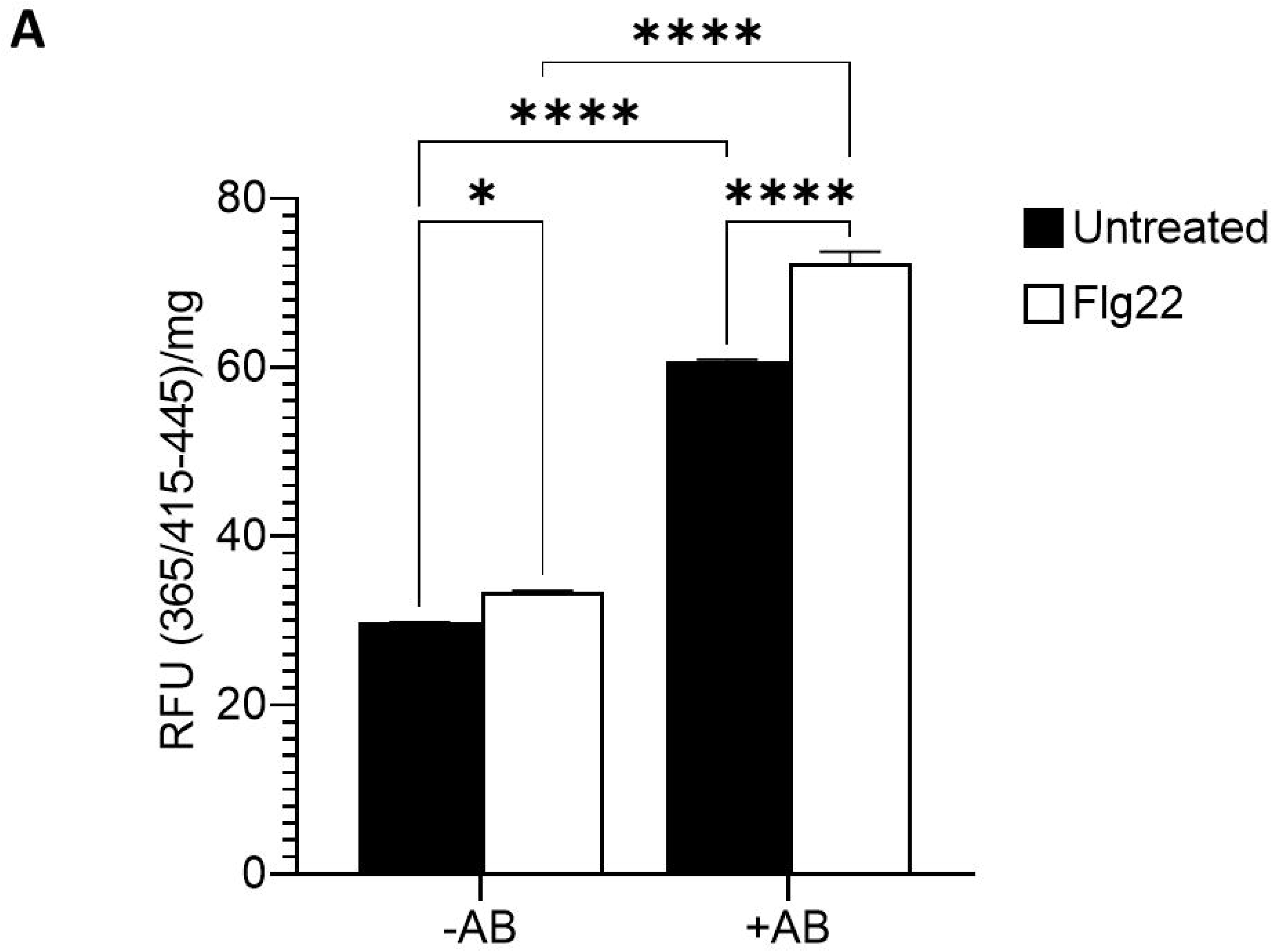
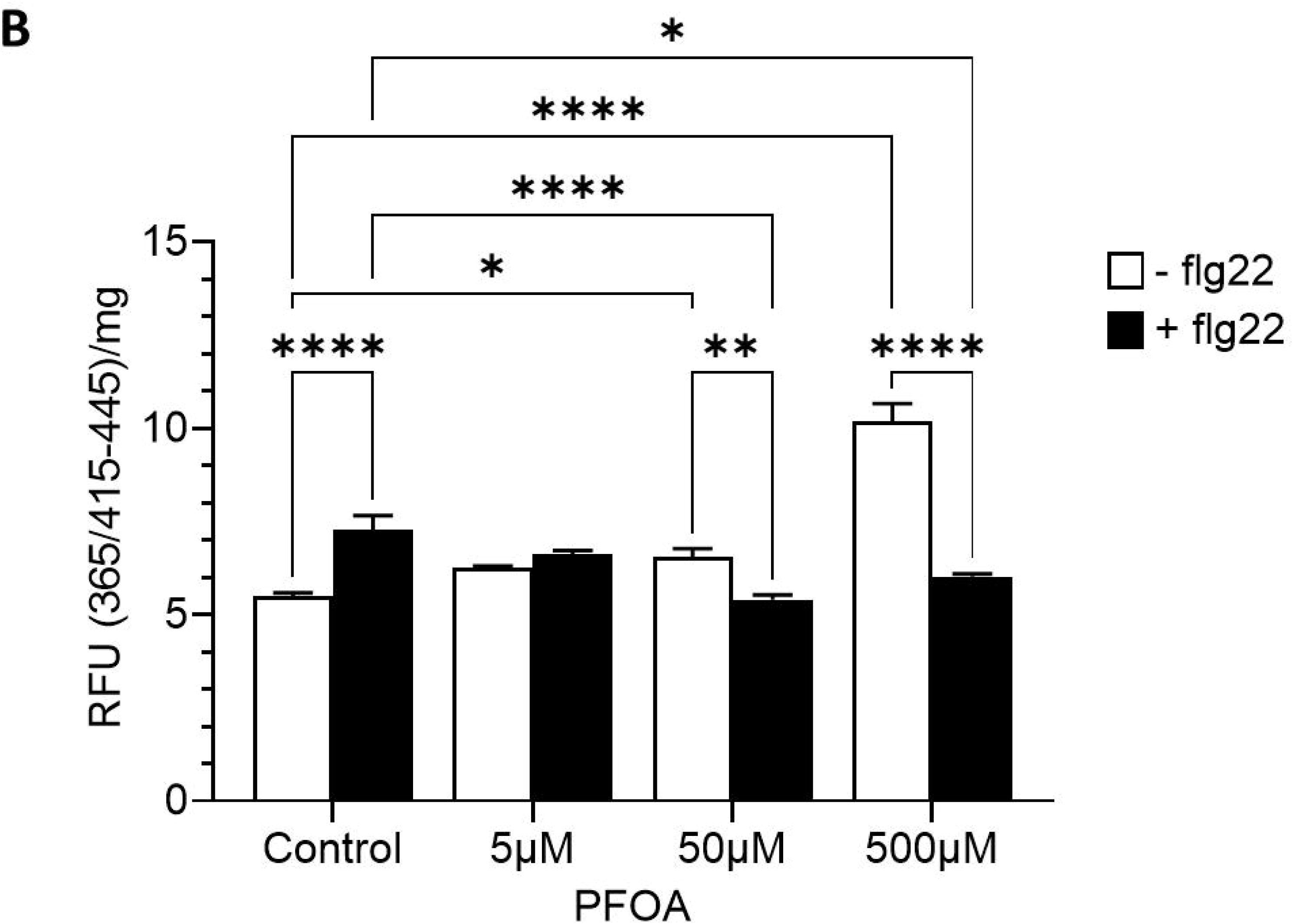
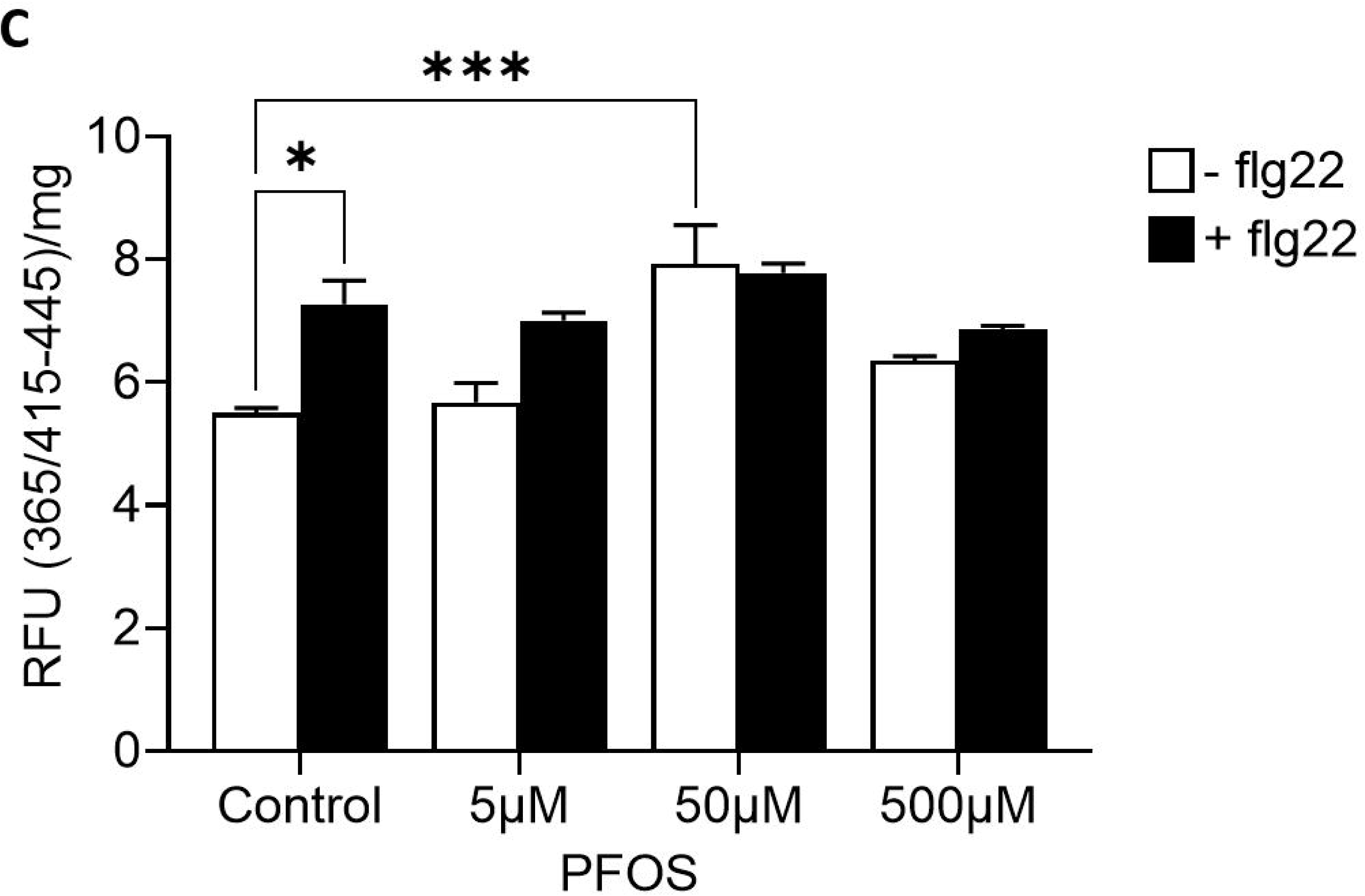
PFAS increases baseline callose production in plants. Mung bean seeds were germinated in untreated conditions (**A**) or in the presence of PFAS (**B, C**) and allowed to grow for 29 days. As control (**A**), true leaves were removed, weighed, challenged with Flg22 (+Flg22), stained for callose using Aniline blue (+AB), measured for fluorescence at 365 (ex)/415-445 (em) nm, spectrophotometrically, and normalised against the weight of leaves. For **B and C**, leaves were obtained from mung bean plants grown in the presence of PFOA (**B**) or PFOS (**C**), challenged with Flg22 (+Flg22) and processed as described in the Materials and Methods section. Significance was determined by two-way ANOVA, and a Tukey’s multiple comparisons test. (****) p ≤ 0.0001, (*), p ≤ 0.05. Results are expressed as the mean ± SD of at least two independent experiments (n = 6 leaves/condition).

## DISCUSSION

The toxic effects of PFAS on the biology of mung bean seeds (*Vigna radiata*) is reported in this study. PFAS are notoriously persistent in the environment and is not easily biodegradable. Their uptake by food crops and subsequent transfer into the human food chain raises particular concerns regarding their uptake by food crops and subsequent transfer into the human food chain (**Adu et al., 2023**) Although PFAS is known to affect humans and animals, causing cancer, organ failure and developmental defects (**Ferretti et al., 2026**), our knowledge on its impact on plants is limited. We found a high dose of high concentrations of PFOA delayed germination of *V. radiata* seeds, when compared to PFOS. This is unsurprising, as it has been shown elsewhere that high concentrations of PFOA inhibit germination and seedling development in wheat (*Triticum aestivum* L.), lettuce (*Lactuca sativa*) and pakchoi (*Brassica rapa chinensis*) was negatively impacted by PFOA not PFOS (**Zhou et al, 2016; Li, 2009**). As PFAS is water soluble and hydrophobic, they are able to be taken up by plants via water in soil pores. PFAS, such as PFOA and PFOS, are readily translocated through the root barriers containing lipid root membranes via their hydrophobic, long carbon chain before entering the xylem (**Glüge et al, 2020**). Furthermore, while seeds grown in both PFOA and PFOS conditions eventually germinated, significant defects in <u>root</u>, <u>hypocotyl</u> and <u>leaf</u> development were observed at high dose of PFOA compared to PFOS. Interestingly, this differential response is also observed in other plants, with PFOS preferentially retained within root tissues, while PFOA are more readily translocated to plant parts above-ground, suggesting the difference in functional groups (PFOS, sulfonate; PFOA, carboxylate) and not just chain length, determines distribution within plants (**Knight et al., 2021; Zhi et al., 2022; Xu et al., 2022, 2024**).

Interestingly, potatoes have demonstrated higher levels of PFOA accumulated, compared to PFOS in the tubers whereas the opposite is true of potato peels (**Stahl et al, 2009**). Therefore, our results could be due to differential localisation of PFOA and PFOS in parts of the seeds that leads to short-term success in germination but long-term defects in plant development. Interestingly, within a narrower range of PFAS concentrations (5-500 μM), while there is dose dependent decrease in seedling growth under PFOS conditions (less so in PFOA), the short-term impact is more observable compared to the long-term impact. This is seen in the hypocotyl length while not having impact on seed germination. This result differs from **Li et al (2009)** who demonstrated that seed germination of lettuce (*Lactuca sativa*) and pakchoi (*Brassica rapa chinensis*) was negatively impacted by PFOA, not PFOS. Likewise, the germination of Chinese cabbage and wheat was reduced by PFOA and PFOS contamination, respectively (**De-Yong et al 2011; Zhou et al, 2016**). This suggests that PFAS poisoning is species dependent and there are mechanisms for recovery with potential PFAS poisoning.

The impact on growth – as evidenced from wet (or fresh) weight at the lowest dose (5 μM), is more pronounced with PFOA, not PFOS. However, at higher doses (500 μM), both PFAS have equally deleterious effects on mung bean seedling weights, leaf sizes and leaf weights. Interestingly, both PFAS had no effect on the dry weights of mung bean seedlings. A similar phenotype was observed in *Arabidopsis thaliana* where there was no change with dry weight when exposed to low dose (**2** μ**g/l)** PFOA or PFOS (equivalent to **4.84 nM PFOA, 4 nM PFOS)** (**Müller et al 2015**). Furthermore, **1 mg/l** (equivalent to **2.42** μ**M PFOA,** 2 μ**M PFOS**) of PFOS or PFOA exhibited no dry weight effect on maize growth too. However, at higher doses of **10 mg/l (equivalent to 24.15** μ**M PFOA, 20.00** μ**M PFOS)**, the dry weights of maize roots and shoots decreased significantly (**Wen et al, 2013**).

Recently it was shown that **10 – 100** μ**M of PFOA** led to a significant decrease in the wet weight of lettuce (*Lactuca sativa*) and radish (*Raphanus sativus*), but not carrot (*Daucus carota*) (**Ozden, 2026**). This suggests that the presence of PFOA and PFOS leads to different responses. As the leaf weights and sizes, along with wet weight of mung bean seedlings, fluctuates in the presence of both PFOA and PFOS even though it had minimal impact on the dry weight, this suggest PFAS affects water retention capability. This is because PFAS can be taken up by the root system via aquaporins, ion channels, or carrier proteins (**Goswami et al., 2025; Mei et al., 2021**). Whether PFAS have translocated to the aerial tissue of the mung bean plant, and to what extent, needs further study. But it has been shown that plants can also absorb and sequester contaminants by their roots, thereby immobilising these contaminants and preventing its further spread by water movement – as observed in lead (Pb) trapped in the roots of *Matthiola* (Brassicaceae) species (**Bolan et al., 2011; Salehi-Eskandari et al., 2022**).

Interestingly, PFOA alters water uptake in *Alisma orientale* (Asian water plantain), and changes the structures, compositions and functions of lipids, proteins and DNA, as observed using Fourier transform infrared spectroscopy (FTIR) (**Wang et al., 2022**). Furthermore, *Salix triandra* (Willow) trees exposed to a hydroponic solution containing 11 different PFAS increased water retention in roots (not leaves), reduced stomatal conductance (smaller stomata, or leaf pores, openings) and increased xylem vulnerability to cavitation, or the formation of air bubbles that block water flow and increases the risk of water transport failure (**Battisti et al., 2023**). Therefore, as plants maintains water homeostasis by balancing water uptake from soil with transpiration losses, there is a need for greater study into the actual molecular and cellular impact that PFAS have on overall plant health and function.

Chlorophyll which is vital to plant growth and crop production also showed variability in response to PFAS species and dose. In mung beans leaves, chlorophyll levels increased in the presence of PFAS, more so with low dose PFOA than PFOS. This may seem surprising as chlorophyll levels were reduced and photosynthesis functions impaired in plants exposed to PFAS e.g. *Cucumis sativus* (cucumber) and *Arabidopsis thaliana* (thale cress) and *Triticum aestivum* L. (wheat), (**Fan et al., 2020; Du et al., 2020; Lin et al., 2020**). On the other hand, species like *Cucurbita pepo* (field pumpkin) and *Daucus carota* (carrot) had an increase in chlorophyll levels when exposed to a mixture of PFAS or 100 µM of PFOA, respectively (**XXX**; **Ozden, 2026**). Some species, like *Brassica juncea* (mustard greens), showed tolerance and little change to chlorophyll levels upon exposure to PFAS (**Wattier et al., 2023**). Therefore, the overall picture shows that PFAS does have an impact on chlorophyll concentration, though this dose and both plant and chemical species dependent.

Finally, we explored the impact PFAS might have on a plant’s stress response. While PFAS is known to exacerbate oxidative stress in mammalian cells by raising malondialdehyde (MDA) and reactive oxygen species (ROS) levels and lowering antioxidant enzyme capacity in aquatic and terrestrial organisms (**Chen et al., 2023; Bonato et al., 2020; Panaretakis et al., 2001; Qian et al., 2010; Reistad et al., 2013**). Plants responses to stress are different from animals and one of them is the accumulation of callose, a polysaccharide produced by callose synthases, which exists in the cell walls of a wide variety of plants and detectable by aniline blue (**Wang et al., 2022; Schenk and Schikora, 2015**). It plays important roles during plant development and/or in response to stress (**Chen and Kim, 2009**). Therefore, it would be important to understand if PFOA and/or PFOS are able to induce a stress response in mung bean seedlings, through the production of callose. Mung bean leaves produced callose upon treatment with flagellin-22 fragment ex vivo, as expected, as bacterial flagellin is an effective elicitor in both plants and algae, and is part of the Pattern-Triggered Immunity (PTI) (**Schenk and Schikora, 2015; Schenk et al., 2014; Jones et al., 2024**). However, while there was an increase in callose production in plants exposed to PFOA at high dose or PFOS at medium dose, this elevated stress response was not further increased when challenged by flagellin-22. This suggests that plant immunity may be primed by PFAS which prevents further infection. This is an interesting piece of research as environmental contaminants such as ozone or heavy metals are known to either generate reactive oxygen species (ROS) which perturbs leaf’s defence enzymes or disrupt enzymatic antioxidant defences inside leaf tissues, respectively, thereby weakening overall resistance (**Jones et al., 2024)**. But low dose of environmental contaminants – and other triggers such as microorganisms or chemical agents – can prime the immune system in plants e.g. heavy metals in tomatoes, in a process known as “induced resistance” (**Chakraborty et al., 2015; Flors et al., 2024**). It is likely that low dose PFAS has the ability of inducing resistance in plants, and further work is required to explore this phenomenon.

## ACKNOWLEDGEMENTS

JL would like to thank his family for their help and support and to Zarah Pattison for her support with editing and proof-reading of the manuscript. The University of Stirling and the Carnegie Trust for the Universities of Scotland (RIG008296) provided some financial support.

## DECLARATION OF GENERATIVE AI AND AI-ASSISTED TECHNOLOGIES IN THE MANUSCRIPT PREPARATION PROCESS

During the preparation of this work the author used jenni.ai and ChatGPT in order to prepare parts of the Introduction and the Discussion. After using this tool/service, the author(s) reviewed and edited the content as needed and take(s) full responsibility for the content of the published article.

**Supplementary Figure:** Absorption spectrum of *Vigna radiata* leaf extract. Control (untreated) leaves from 29-day old mung bean plants were suspended in hydroponic water, chlorophyll extracted and scanned spectrophotometrically from 350-900 nm, as described in the Materials and Methods section.

## REFERENCES

1. Adu O, Ma X, Sharma VK. Bioavailability, phytotoxicity and plant uptake of per-and polyfluoroalkyl substances (PFAS): A review. Journal of Hazardous Materials 2023;447:130805. 10.1016/j.jhazmat.2023.130805.

2. Battisti I, Zambonini D, Ebinezer LB et al. Perfluoroalkyl substances exposure alters stomatal opening and xylem hydraulics in willow plants. Chemosphere 2023;344:140380. 10.1016/j.chemosphere.2023.140380.

3. Biswas B, Joseph A, Parveen N et al. Contamination of per- and poly-fluoroalkyl substances in agricultural soils: A review. Journal of Environmental Management 2025;380:124993. 10.1016/j.jenvman.2025.124993.

4. Bolan NS, Park JH, Robinson B et al. Phytostabilization. In: Advances in Agronomy. n.p.: Elsevier, 2011, 112.145–204. 10.1016/B978-0-12-385538-1.00004-4.

5. Bonato M, Corrà F, Bellio M et al. PFAS Environmental Pollution and Antioxidant Responses: An Overview of the Impact on Human Field. IJERPH 2020;17(21):8020. 10.3390/ijerph17218020.

6. Caniglia J, Snow DD, Messer T et al. Extraction, analysis, and occurrence of per- and polyfluoroalkyl substances (PFAS) in wastewater and after municipal biosolids land application to determine agricultural loading. Front Water 2022;4:892451. 10.3389/frwa.2022.892451.

7. Chakraborty N, Chandra S, Acharya K. Sublethal Heavy Metal Stress Stimulates Innate Immunity in Tomato. The Scientific World Journal 2015;2015(1):208649. 10.1155/2015/208649.

8. Chen XY, Kim JY. Callose synthesis in higher plants. Plant Signaling & Behavior 2009;4(6):489–92. 10.4161/psb.4.6.8359.

9. Chen JC, Baumert BO, Li Y et al. Associations of per- and polyfluoroalkyl substances, polychlorinated biphenyls, organochlorine pesticides, and polybrominated diphenyl ethers with oxidative stress markers: A systematic review and meta-analysis. Environmental Research 2023;239:117308. 10.1016/j.envres.2023.117308.

10. Costello MCS, Lee LS. Sources, Fate, and Plant Uptake in Agricultural Systems of Per- and Polyfluoroalkyl Substances. Curr Pollution Rep 2020;10(4):799–819. 10.1007/s40726-020-00168-y.

11. De-Yong Z, Xiu-Ying S, Xiao-Lu X. The effects of perfluorooctane sulfonate (PFOS) on Chinese cabbage (Brassica rapa pekinensis) germination and development. Procedia Engineering 2011;18:206–13. 10.1016/j.proeng.2011.11.033.

12. Dimitrakopoulou ME, Karvounis M, Marinos G et al. Comprehensive analysis of PFAS presence from environment to plate. Npj Sci Food 2024;8(1):80. 10.1038/s41538-024-00319-1.

13. Du W, Liu X, Zhao L et al. Response of cucumber (Cucumis sativus) to perfluorooctanoic acid in photosynthesis and metabolomics. Science of The Total Environment 2020;724:138257. 10.1016/j.scitotenv.2020.138257.

14. Ecke F, Ytrehus B, Evander M et al. Biomagnification and potential health effects of per- and polyfluoroalkyl substances (PFAS) in a terrestrial food web. Sci Rep 2025;15(1):31003. 10.1038/s41598-025-16395-6.

15. Everwand G, Cass S, Dauber J et al. Legume crops and biodiversity. In: Murphy-Bokern D, Stoddard FL, Watson CA (eds), Legumes in Cropping Systems, 1st edn. UK: CABI, 2017, 55–69. 10.1079/9781780644981.0055.

16. Evich MG, Davis MJB, McCord JP et al. Per- and polyfluoroalkyl substances in the environment. Science 2022;375(6580):eabg9065. 10.1126/science.abg9065.

17. Fan L, Tang J, Zhang D et al. Investigations on the phytotoxicity of perfluorooctanoic acid in Arabidopsis thaliana. Environ Sci Pollut Res 2020;27(1):1131–43. 10.1007/s11356-019-07018-5.

18. Ferretti F, Barbarossa A, Bardhi A. An overview of the impact of PFAS on animals, humans, and the environment using a One Health approach. Environ Sci Pollut Res 2026;33(5):1461–89. 10.1007/s11356-026-37412-9.

19. Ferreira H, Pinto E, Vasconcelos MW. Legumes as a Cornerstone of the Transition Toward More Sustainable Agri-Food Systems and Diets in Europe. Front Sustain Food Syst 2021;5:694121. 10.3389/fsufs.2021.694121.

20. Flors V, Kyndt T, Mauch-Mani B et al. Enabling sustainable crop protection with induced resistance in plants. Front Sci 2024;2:1407410. 10.3389/fsci.2024.1407410.

21. Glüge J, Scheringer M, Cousins IT et al. An overview of the uses of per- and polyfluoroalkyl substances (PFAS). Environ Sci: Processes Impacts 2020;22(12):2345–73. 10.1039/D0EM00291G.

22. Goswami L, Dikshit PK, Prakash A et al. A critical review on occurrence, speciation, mobilization, and toxicity of per- and polyfluoroalkyl substances in the soil-microbe-plant system and bioremediation strategies. Journal of Hazardous Materials 2025;494:138743. 10.1016/j.jhazmat.2025.138743.

23. Han F, Zhang S, Zhou W et al. Fabrication and characterization of Pickering high internal phase emulsion stabilized by mung bean flour. Food Sci Technol 2022;42:e85122. 10.1590/fst.85122.

24. Hannachi S, Signore A, Adnan M et al. Single and Associated Effects of Drought and Heat Stresses on Physiological, Biochemical and Antioxidant Machinery of Four Eggplant Cultivars. Plants 2022;11(18):2404. 10.3390/plants11182404.

25. Huang W, Ratkowsky DA, Hui C et al. Leaf Fresh Weight Versus Dry Weight: Which is Better for Describing the Scaling Relationship between Leaf Biomass and Leaf Area for Broad-Leaved Plants? Forests 2019;10(3):256. 10.3390/f10030256.

26. Idris OA, Erasmus M. Degradation pathways of perfluoroalkyl and polyfluoroalkyl compounds: Removal in water and soil using fungi and plant-based remediation. Environmental Advances 2024;18:100598. 10.1016/j.envadv.2024.100598.

27. Ievinsh G. Water Content of Plant Tissues: So Simple That Almost Forgotten? Plants 2023;12(6):1238. 10.3390/plants12061238.

28. Islam Md Anwarul, Parvin MI, Nguyen C et al. Per- and polyfluoroalkyl substances (PFAS) contamination in agriculture and its potential conflict with circular economy. Environmental Pollution 2025;385:127036. 10.1016/j.envpol.2025.127036.

29. Islam Mohammad Aminul. 112 PUBLICATIONS 2,181 CITATIONS SEE PROFILE. J Anim Plant Sci n.d.

30. Jha G, Kankarla V, McLennon E et al. Per- and Polyfluoroalkyl Substances (PFAS) in Integrated Crop–Livestock Systems: Environmental Exposure and Human Health Risks. IJERPH 2021;18(23):12550. 10.3390/ijerph182312550.

31. Jones JDG, Staskawicz BJ, Dangl JL. The plant immune system: From discovery to deployment. Cell 2024;187(9):2095–116. 10.1016/j.cell.2024.03.045.

32. Kapravelou G, Martínez R, Perazzoli G et al. Germination Improves the Polyphenolic Profile and Functional Value of Mung Bean (Vigna radiata L.). Antioxidants 2020;9(8):746. 10.3390/antiox9080746.

33. Khalil N, Ducatman AM, Sinari S et al. Per- and Polyfluoroalkyl Substance and Cardio Metabolic Markers in Firefighters. Journal of Occupational & Environmental Medicine 2020;62(12):1076–81. 10.1097/JOM.0000000000002062.

34. Kohler A, Schwindling S, Conrath U. Extraction and Quantitative Determination of Callose from *Arabidopsis* Leaves. BioTechniques 2000;28(6):1084–6. 10.2144/00286bm06.

35. Knight ER, Bräunig J, Janik LJ et al. An investigation into the long-term binding and uptake of PFOS, PFOA and PFHxS in soil – plant systems. Journal of Hazardous Materials 2021;404:124065. 10.1016/j.jhazmat.2020.124065.

36. Kumar D, Singh H, Raj S et al. Chlorophyll a fluorescence kinetics of mung bean (Vigna radiata L.) grown under artificial continuous light. Biochemistry and Biophysics Reports 2020;24:100813. 10.1016/j.bbrep.2020.100813.

37. Lankin VZ, Tikhaze AK, Melkumyants AM. Malondialdehyde as an Important Key Factor of Molecular Mechanisms of Vascular Wall Damage under Heart Diseases Development. IJMS 2022;24(1):128. 10.3390/ijms24010128.

38. Lenka SP, Kah M, Padhye LP. A review of the occurrence, transformation, and removal of poly- and perfluoroalkyl substances (PFAS) in wastewater treatment plants. Water Research 2021;199:117187. 10.1016/j.watres.2021.117187.

39. Lesmeister L, Lange FT, Breuer J et al. Extending the knowledge about PFAS bioaccumulation factors for agricultural plants – A review. Science of The Total Environment 2021;766:142640. 10.1016/j.scitotenv.2020.142640.

40. Li M. Toxicity of perfluorooctane sulfonate and perfluorooctanoic acid to plants and aquatic invertebrates. Environmental Toxicology 2009;24(1):95–101. 10.1002/tox.20396.

41. Li FM, Lu ZG, Yue M. Analysis of Photosynthetic Characteristics and UV-B Absorbing Compounds in Mung Bean Using UV-B and Red LED Radiation. Journal of Analytical Methods in Chemistry 2014;2014:1–5. 10.1155/2014/378242.

42. Li L, Yang T, Liu R et al. Food legume production in China. The Crop Journal 2017;5(2):115–26. 10.1016/j.cj.2016.06.001.

43. Li J, Sun J, Li P. Exposure routes, bioaccumulation and toxic effects of per- and polyfluoroalkyl substances (PFASs) on plants: A critical review. Environment International 2022;158:106891. 10.1016/j.envint.2021.106891.

44. Lin Q, Zhou C, Chen L et al. Accumulation and associated phytotoxicity of novel chlorinated polyfluorinated ether sulfonate in wheat seedlings. Chemosphere 2020;249:126447. 10.1016/j.chemosphere.2020.126447.

45. Luche S, Buczny J, Visioli G. Plant biomass responses to PFAS exposure: A meta-analysis with implications for phytoremediation in terrestrial and aquatic systems. Journal of Environmental Management 2026;403:129217. 10.1016/j.jenvman.2026.129217.

46. Luche S, Fanelli A, Buschini A et al. PFOA and PFOS accumulation induces genotoxic damage and proteomic alterations in Cannabis sativa shoots. Journal of Hazardous Materials 2026;501:140686. 10.1016/j.jhazmat.2025.140686.

47. Mao W, Li M, Xue X et al. Bioaccumulation and toxicity of perfluorooctanoic acid and perfluorooctane sulfonate in marine algae Chlorella sp. Science of The Total Environment 2023;870:161882. 10.1016/j.scitotenv.2023.161882.

48. Marzi D, Valente F, Luche S et al. Phytoremediation of perfluoroalkyl and polyfluoroalkyl substances (PFAS): Insights on plant uptake, omics analysis, contaminant detection and biomass disposal. Science of The Total Environment 2025;959:178323. 10.1016/j.scitotenv.2024.178323.

49. Mei W, Sun H, Song M et al. Per- and polyfluoroalkyl substances (PFASs) in the soil– plant system: Sorption, root uptake, and translocation. Environment International 2021;156:106642. 10.1016/j.envint.2021.106642.

50. Mei W, Sun H, Song M et al. Per- and polyfluoroalkyl substances (PFASs) in the soil– plant system: Sorption, root uptake, and translocation. Environment International 2021;156:106642. 10.1016/j.envint.2021.106642.

51. Mei W, Sun H, Song M et al. Per- and polyfluoroalkyl substances (PFASs) in the soil– plant system: Sorption, root uptake, and translocation. Environment International 2021;156:106642. 10.1016/j.envint.2021.106642.

52. Menon R. Water Relations in Plants: Physiological Mechanisms and Adaptive Responses to Hydric Stress. 2025.

53. Michalak A, Małas K, Dąbrowska K et al. Molecular Mechanisms and Experimental Strategies for Understanding Plant Drought Response. Plants 2026;15(1):149. 10.3390/plants15010149.

54. Mulabagal V, Liu L, Qi J et al. A rapid UHPLC-MS/MS method for simultaneous quantitation of 23 perfluoroalkyl substances (PFAS) in estuarine water. Talanta 2018;190:95–102. 10.1016/j.talanta.2018.07.053.

55. Müller CE, LeFevre GH, Timofte AE et al. Competing mechanisms for perfluoroalkyl acid accumulation in plants revealed using an *Arabidopsis* model system. Environmental Toxicology and Chemistry 2015;35(5):1138–47. 10.1002/etc.3251.

56. Ofoegbu PC, Wagner DC, Abolade O et al. Impacts of perfluorooctanesulfonic acid on plant biometrics and grain metabolomics of wheat (Triticum aestivum L.). Journal of Hazardous Materials Advances 2022;7:100131. 10.1016/j.hazadv.2022.100131.

57. Okpanachi IY, Gotan YS, Dauda S et al. Combined effect of PFOS and PFOA on phytoplankton interactions and overall community health. Journal of Plankton Research 2025;47(5):fbaf044. 10.1093/plankt/fbaf044.

58. Omagamre EW, Mansourian Y, Liles D et al. Perfluorobutanoic Acid (PFBA) Induces a Non-Enzymatic Oxidative Stress Response in Soybean (Glycine max L. Merr.). IJMS 2022;23(17):9934. 10.3390/ijms23179934.

59. Oviedo-Vargas D, Anton J, Coleman-Kammula S et al. Quantification of PFAS in soils treated with biosolids in ten northeastern US farms. Sci Rep 2025;15(1):5582. 10.1038/s41598-025-90184-z.

60. Ozden H. The effects of perfluorooctanoic acid (PFOA) on physiological processes and oxidative damage in edible vegetables. Open Life Sciences 2026;21(1):20251250. 10.1515/biol-2025-1250.

61. Pan CG, Sun RX. Understanding PFAS: Occurrence, Fate, Removal, and Effects. Toxics 2024;12(8):605. 10.3390/toxics12080605.

62. Panaretakis T, Shabalina IG, Grandér D et al. Reactive Oxygen Species and Mitochondria Mediate the Induction of Apoptosis in Human Hepatoma HepG2 Cells by the Rodent Peroxisome Proliferator and Hepatocarcinogen, Perfluorooctanoic Acid. Toxicology and Applied Pharmacology 2001;173(1):56–64. 10.1006/taap.2001.9159.

63. Paulson SG, Liu S, Rotty JD. Physical confinement and phagocytic uptake induce persistent cell migration. Biology Open 2025;14(9):bio062021. 10.1242/bio.062021.

64. Pavlovic D, Nikolic B, Djurovic S et al. Chlorophyll as a measure of plant health: Agroecological aspects. Pesticidi i Fitomedicina 2014;29(1):21–34. 10.2298/PIF1401021P.

65. Peñas E, Gómez R, Frías J et al. Effects of combined treatments of high pressure, temperature and antimicrobial products on germination of mung bean seeds and microbial quality of sprouts. Food Control 2010;21(1):82–8. 10.1016/j.foodcont.2009.04.008.

66. Peritore AF, Gugliandolo E, Cuzzocrea S et al. Current Review of Increasing Animal Health Threat of Per- and Polyfluoroalkyl Substances (PFAS): Harms, Limitations, and Alternatives to Manage Their Toxicity. IJMS 2023;24(14):11707. 10.3390/ijms241411707.

67. Pietrini F, Wyrwicka-Drewniak A, Passatore L et al. PFOA accumulation in the leaves of basil (Ocimum basilicum L.) and its effects on plant growth, oxidative status, and photosynthetic performance. BMC Plant Biol 2024;24(1):556. 10.1186/s12870-024-05269-0.

68. Qian Y, Ducatman A, Ward R et al. Perfluorooctane Sulfonate (PFOS) Induces Reactive Oxygen Species (ROS) Production in Human Microvascular Endothelial Cells: Role in Endothelial Permeability. Journal of Toxicology and Environmental Health, Part A 2010;73(12):819–36. 10.1080/15287391003689317.

69. Ranawake A, Dahanayaka N, Amarasingha U et al. Effect of Water Stress on Growth and Yield of Mung Bean (Vigna radiata L). Trop Agric Res & Ext 2012;14(4). 10.4038/tare.v14i4.4851.

70. Reistad T, Fonnum F, Mariussen E. Perfluoroalkylated compounds induce cell death and formation of reactive oxygen species in cultured cerebellar granule cells. Toxicology Letters 2013;218(1):56–60. 10.1016/j.toxlet.2013.01.006.

71. Salehi-Eskandari B, Gahrouei MS, Boyd RS et al. Physiological responses to lead and PEG-simulated drought stress in metallicolous and non-metallicolous Matthiola (Brassicaceae) species from Iran. South African Journal of Botany 2022;150:1011–21. 10.1016/j.sajb.2022.09.016.

72. Sarma RK, Saikia R. Alleviation of drought stress in mung bean by strain Pseudomonas aeruginosa GGRJ21. Plant Soil 2014;377(1–2):111–26. 10.1007/s11104-013-1981-9.

73. Schenk ST, Hernández-Reyes C, Samans B et al. *N* -Acyl-Homoserine Lactone Primes Plants for Cell Wall Reinforcement and Induces Resistance to Bacterial Pathogens via the Salicylic Acid/Oxylipin Pathway. Plant Cell 2014;26(6):2708–23. 10.1105/tpc.114.126763.

74. Schenk S, Schikora A. Staining of Callose Depositions in Root and Leaf Tissues. BIO-PROTOCOL 2015;5(6). 10.21769/BioProtoc.1429.

75. Sparks DL. Advances in Agronomy. Amsterdam: Elsevier, 2011.

76. Stahl T, Heyn J, Thiele H et al. Carryover of Perfluorooctanoic Acid (PFOA) and Perfluorooctane Sulfonate (PFOS) from Soil to Plants. Arch Environ Contam Toxicol 2009;57(2):289–98. 10.1007/s00244-008-9272-9.

77. Tarnocai D, Geo P, McDonald L et al. ›Comparison of Site Specific and Literature Bioconcentration Factors and Implications for Site Specific Target Level Derivation. n.d.

78. Valente F, Chisini Granzotto G, Panozzo A et al. PFAS-induced morpho-physiological, photosynthetic and tissue accumulation responses of pot-grown hemp, sunflower and maize under soil amendment with humic acids. Front Plant Sci 2026;17:1791472. 10.3389/fpls.2026.1791472.

79. Van Grondelle R, Boeker E. Limits on Natural Photosynthesis. J Phys Chem B 2017;121(30):7229–34. 10.1021/acs.jpcb.7b03024.

80. Voijant Tangahu B. Growth Rate Measurement of Scirpus Grossus Plant as Preliminary Step to Apply the Plant in Wastewater Treatment Using Reedbed System. J Civil Environ Eng 2016;05(06). 10.4172/2165-784X.1000192.

81. Wang L, Liu Y, Kaur M et al. Phytotoxic Effects of Polyethylene Microplastics on the Growth of Food Crops Soybean (Glycine max) and Mung Bean (Vigna radiata). IJERPH 2021;18(20):10629. 10.3390/ijerph182010629.

82. Wang TT, Wang S, Shao S et al. Perfluorooctanoic acid (PFOA)-induced alterations of biomolecules in the wetland plant Alisma orientale. Science of The Total Environment 2022;820:153302. 10.1016/j.scitotenv.2022.153302.

83. Wang TT, Ying GG, Shi WJ et al. Uptake and Translocation of Perfluorooctanoic Acid (PFOA) and Perfluorooctanesulfonic Acid (PFOS) by Wetland Plants: Tissue- and Cell-Level Distribution Visualization with Desorption Electrospray Ionization Mass Spectrometry (DESI-MS) and Transmission Electron Microscopy Equipped with Energy-Dispersive Spectroscopy (TEM-EDS). Environ Sci Technol 2020;54(10):6009–20. 10.1021/acs.est.9b05160.

84. Wang W, Rhodes G, Ge J et al. Uptake and accumulation of per- and polyfluoroalkyl substances in plants. Chemosphere 2020;261:127584. 10.1016/j.chemosphere.2020.127584.

85. Wang X, Zhang W, Lamichhane S et al. Effects of physicochemical properties and co-existing zinc agrochemicals on the uptake and phytotoxicity of PFOA and GenX in lettuce. Environ Sci Pollut Res 2023;30(15):43833–42. 10.1007/s11356-023-25435-5.

86. Wang Y, Niu J, Zhang L et al. Toxicity assessment of perfluorinated carboxylic acids (PFCAs) towards the rotifer Brachionus calyciflorus. Science of The Total Environment 2014;491–492:266–70. 10.1016/j.scitotenv.2014.02.028.

87. Wattier BD, Gonzales AK, Martinez NE. Perfluorooctanoic acid uptake in the mustard species *Brassica juncea*. J of Env Quality 2023;52(1):199–206. 10.1002/jeq2.20431.

88. Wee SY, Aris AZ. Environmental impacts, exposure pathways, and health effects of PFOA and PFOS. Ecotoxicology and Environmental Safety 2023;267:115663. 10.1016/j.ecoenv.2023.115663.

89. Wen B, Li L, Liu Y et al. Mechanistic studies of perfluorooctane sulfonate, perfluorooctanoic acid uptake by maize (Zea mays L. cv. TY2). Plant Soil 2013;370(1–2):345–54. 10.1007/s11104-013-1637-9.

90. Wen B, Wu Y, Zhang H et al. The roles of protein and lipid in the accumulation and distribution of perfluorooctane sulfonate (PFOS) and perfluorooctanoate (PFOA) in plants grown in biosolids-amended soils. Environmental Pollution 2016;216:682–8. 10.1016/j.envpol.2016.06.032.

91. Xu B, Qiu W, Du J et al. Translocation, bioaccumulation, and distribution of perfluoroalkyl and polyfluoroalkyl substances (PFASs) in plants. iScience 2022;25(4):104061. 10.1016/j.isci.2022.104061.

92. Xu J, Cui Q, Ren H et al. Differential uptake and translocation of perfluoroalkyl substances by vegetable roots and leaves: Insight into critical influencing factors. Science of The Total Environment 2024;949:175205. 10.1016/j.scitotenv.2024.175205.

93. Yan H, Zhao T, Xu B et al. Arbuscular mycorrhizal fungi suffer from perfluorooctanoic acid in soil but ameliorate negative effects on plant performance. Applied Soil Ecology 2025;212:106169. 10.1016/j.apsoil.2025.106169.

94. Yang X, Ye C, Liu Y et al. Accumulation and phytotoxicity of perfluorooctanoic acid in the model plant species Arabidopsis thaliana. Environmental Pollution 2015;206:560–6. 10.1016/j.envpol.2015.07.050.

95. Zhang J, Naveed H, Chen K et al. Toxicity of Per- and Polyfluoroalkyl Substances and Their Substitutes to Terrestrial and Aquatic Invertebrates—A Review. Toxics 2025;13(1):47. 10.3390/toxics13010047.

96. Zhang L, Niu J, Li Y et al. Evaluating the sub-lethal toxicity of PFOS and PFOA using rotifer Brachionus calyciflorus. Environmental Pollution 2013;180:34–40. 10.1016/j.envpol.2013.04.031.

97. Zhang L, Niu J, Li Y et al. Evaluating the sub-lethal toxicity of PFOS and PFOA using rotifer Brachionus calyciflorus. Environmental Pollution 2013;180:34–40. 10.1016/j.envpol.2013.04.031.

98. Zhang P, Sun L, Liu F et al. Perfluorooctanoic acid and perfluorooctane sulfonic acid inhibit plant growth through the modulation of phytohormone signalling pathways: Evidence from molecular and genetic analysis in Arabidopsis. Science of The Total Environment 2022;851:158287. 10.1016/j.scitotenv.2022.158287.

99. Zhang P, Sun L, Liu F et al. Perfluorooctanoic acid and perfluorooctane sulfonic acid inhibit plant growth through the modulation of phytohormone signalling pathways: Evidence from molecular and genetic analysis in Arabidopsis. Science of The Total Environment 2022;851:158287. 10.1016/j.scitotenv.2022.158287.

100. Zhi Y, Zhao X, Qian S et al. Removing emerging perfluoroalkyl ether acids and fluorotelomer sulfonates from water by nanofiltration membranes: Insights into performance and underlying mechanisms. Separation and Purification Technology 2022;298:121648. 10.1016/j.seppur.2022.121648.

101. Zhou L, Xia M, Wang L et al. Toxic effect of perfluorooctanoic acid (PFOA) on germination and seedling growth of wheat (Triticum aestivum L.). Chemosphere 2016;159:420–5. 10.1016/j.chemosphere.2016.06.045.

