## Supplementary figures and images for "Plant functional defects experienced upon growth under Per-/Poly-fluoroalkyl substances (PFAS) conditions"

### Supplementary Figure:

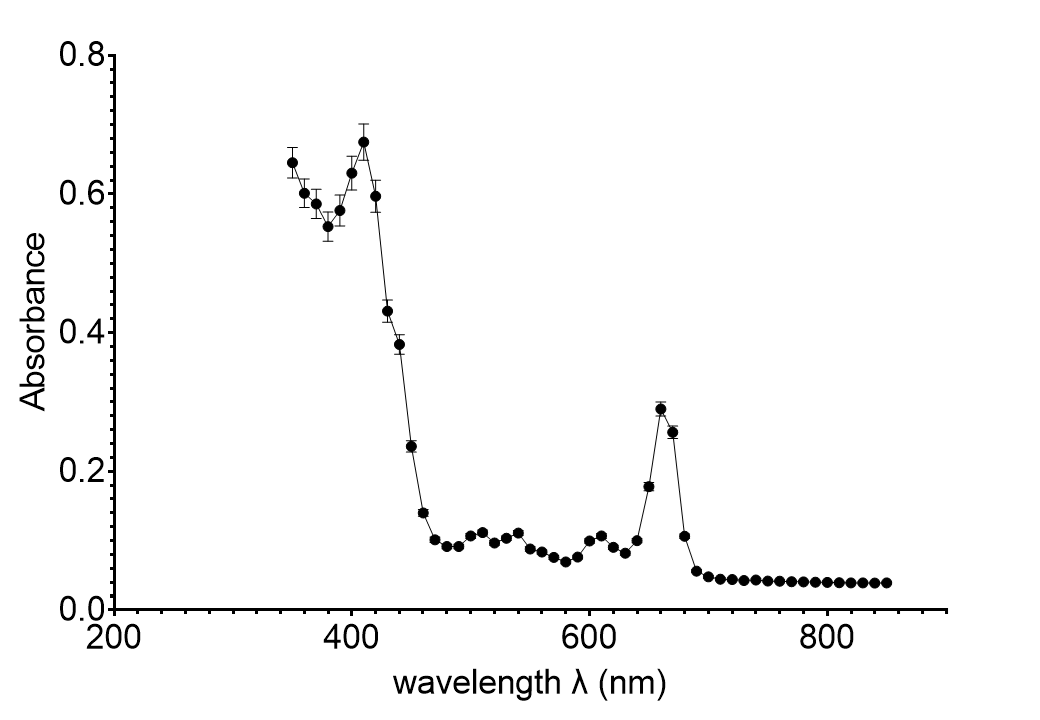
